# Topoisomerase II deficiency leads to a postreplicative structural shift in all *Saccharomyces cerevisiae* chromosomes

**DOI:** 10.1101/2020.09.17.301317

**Authors:** Jessel Ayra-Plasencia, Cristina Ramos-Pérez, Silvia Santana-Sosa, Oliver Quevedo, Sara Medina-Suarez, Emiliano Matos-Perdomo, Marcos Zamora-Dorta, Grant W Brown, Michael Lisby, Félix Machín

**Affiliations:** Unidad de Investigación, Hospital Universitario Nuestra Señora de Candelaria, Tenerife, Spain; Escuela de Doctorado y Estudios de Postgrado, Universidad de La Laguna, Tenerife, Spain; Department of Biochemistry and Donnelly Centre, University of Toronto, Canada; Department of Biology, University of Copenhagen, Denmark; Instituto de Tecnologías Biomédicas, Universidad de la Laguna, Tenerife, Spain; Facultad de Ciencias de la Salud, Universidad Fernando Pessoa Canarias, Las Palmas de Gran Canaria, Spain

**Keywords:** Top2, replication, recombination, Pulsed-Field Gel Electrophoresis (PFGE), ribosomal DNA (rDNA)

## Abstract

The key role of Topoisomerase II (Top2) is the removal of topological intertwines between sister chromatids. In yeast, inactivation of Top2 brings about distinct cell cycle responses. In the case of the conditional *top2-5* allele, interphase and mitosis progress on schedule but cells suffer from a segregation catastrophe. We here show that *top2-5* chromosomes fail to enter a Pulsed-Field Gel Electrophoresis (PFGE) in the first cell cycle, a behavior traditionally linked to the presence of replication and recombination intermediates. We distinguished two classes of affected chromosomes: the rDNA-bearing chromosome XII, which fails to enter a PFGE at the beginning of S-phase, and all the other chromosomes, which fail at a postreplicative stage. In synchronously cycling cells, this late PFGE retention is observed in anaphase; however, we demonstrate that this behavior is independent of cytokinesis, stabilization of anaphase bridges, spindle pulling forces and even anaphase onset. Strikingly, once the PFGE retention has occurred it becomes refractory to Top2 re-activation. DNA combing, two-dimensional electrophoresis, genetic analyses and GFP-tagged DNA damage markers suggest that non-recombinational modifications of late replication intermediates may account for the shift in the PFGE behavior. The fact that this shift does not trigger G_2_/M checkpoints further supports this statement since checkpoints are active for other replicative stresses in the absence of Top2. We propose that the prolonged absence of Top2 activity leads to a general chromosome structural change. This change might interlock chromatids together with catenations and thus contribute to the formation of anaphase bridges in *top2* mutants.

## INTRODUCTION

Among the physical impediments that preclude sister chromatid segregation in anaphase there are topological intertwinings (catenanes), unfinished replication and unresolved recombination intermediates. The presence of any of these structures gives rise to anaphase bridges that can seriously compromise the genome integrity of the immediate cell lineage [1–3]. Surprisingly, cells from most organisms apparently lack specialised checkpoints to detect these aberrant structures and stop anaphase onset. Rather, they rely on indirect ways to supervise putative segregation problems ahead. For instance, during DNA replication, cells check that the replication fork (RF) does not get stalled or blocked, or that long stretches of single-stranded DNA (ssDNA) are not left behind the RF, yet cells can enter anaphase with unfinished replication if it proceeds too slowly compared to the cell division rate [1,4,5]. Likewise, cells monitor both DNA double-strand breaks (DSBs) and ssDNA during DNA damage and early steps of its repair through the homologous recombination (HR) pathway, but not the direct presence of recombination intermediates that connect the damaged DNA with its sister template [6–8]. Catenations also appear invisible to cell cycle checkpoints, although there still exist controversy about putative G_2_/M checkpoint(s) that sense these topological problems in higher eukaryotes [1,9–11].

Eukaryotic type II topoisomerases (topo II/Top2) are exclusive in removing double-strand DNA (dsDNA) catenations [10,12]. The yeast *Saccharomyces cerevisiae* has been extensively used as a model organism where to assess the physiological roles of Top2. This has been facilitated by the simplicity of this cell model and its genetic engineering, the presence of just one *TOP2* gene, and the ability to generate simple conditional alleles; e.g., *top2* thermosensitive (ts) alleles. Pioneering work in David Botstein’s and Rolf Sternglanz’s labs showed that *top2-ts* yeast cells died as a consequence of passing through anaphase [13,14]. Later work demonstrated that Top2 was needed to avoid anaphase bridges, and that completion of cytokinesis had a major role in killing the cell progeny as it severs these anaphase bridges [15–17]. While the presence of anaphase bridges comprising sister chromatid intertwinings is undisputed, several works have suggested the presence of other linkages that could contribute to the sister chromatid segregation defects in *top2-ts*. Thus, unfinished replication has been observed and mapped to chromosome fragile sites, likely coincident with replication termini sites [18]. Replication defects have also been seen at the ribosomal DNA array (rDNA), especially when Top2 deficiency is combined with other mutations that affect rDNA metabolism [19]. The rDNA locus, located on the chromosome XII right arm, is known to be unique because of its unidirectional replication mechanism, the presence of genetically-programmed RF blocks (RFBs), being highly transcribed by RNA polymerases I and III while mostly epigenetically silent for RNA polymerase II, and for being hyper-recombinogenic and, consequently, expected to present more recombination intermediates than other chromosome regions [2,20].

Confounding matters, not all *top2-ts* alleles bring about the same phenotypes. Whereas most of them (but not all) allow a timely anaphase onset, progression beyond this point is more variable, particularly the degree of cytokinetic completion [14,15,21,22]. The underlying reasons behind these differences are somewhat elusive, although features such as residual Top2-ts activity, the genetic background, and the capability to activate a checkpoint that transiently blocks cytokinesis (NoCut/Abscission checkpoint) might be responsible [22]. In previous works we showed that *top2-5* cells were excellent in both synchronously entering anaphase and quickly severing anaphase bridges through cytokinetic furrow ingression [15,23]. In this report, we have studied in more detail the *top2-5* cell cycle and found that all chromosomes fail separation by pulsed-field gel electrophoresis (PFGE). Except for the rDNA-bearing chromosome XII, chromosomes stop entering the PFG not at early S-phase but in mitosis. The use of mutants and mitotic drugs showed that the PFGE behaviour is independent of HR, spindle forces and even anaphase onset. We propose that Top2 deficiency (*top2-5*) gives rise to substantial chromosome structural changes that cannot however trigger an efficient G_2_/M DNA damage checkpoint. Finally, we show that Top2 actions must take place at the correct time, after which the change in chromosome structure becomes refractory to Top2 activity.

## RESULTS

### Yeast chromosomes stop entering a pulsed-field electrophoretic gel in *top2-5*

In a previous work, we used genetically-modified *TOP2* and *top2-5* ts strains to analyse their first cell cycle by both population and single-cell fluorescence microscopy [15]. Specifically, we GFP-tagged the histone H2A2 (*HTA2* gene) in *bar1* derivatives of the original strains in order to synchronously follow the nuclear cell cycle. We started this work by adding flow cytometry (FACS) and Pulsed Field Gel Electrophoresis (PFGE) to the microscopic analysis. Whereas fluorescence microscopy assesses cell and nuclear morphology as markers of cell cycle progression and nuclear segregation, FACS allows determination of both bulk DNA replication and of the degree of uneven segregation after cytokinesis. In addition, PFGE, in conjunction with Southern blot analysis, gives insights into the structural integrity of individual chromosomes; i.e, intact vs broken chromosomes, gross chromosomal rearrangements, and presence of DNA-mediated linkages [7,24–26]. In the case of the latter scenario, affected chromosomes get trapped in the wells of the PFG.

Both strains were arrested in G_1_ at permissive temperature (25 °C) for 3 h before being released to 37 °C for 4 h. Every 30’, samples were collected for analysis by microscopy, FACS and PFGE. As reported before, fluorescence microscopy showed that the *TOP2* strain proceeded into a normal cell cycle (Figure 1a, left panel); S-phase entry (budding) occurred between 30’-90’ and anaphase onset (H2A2-GFP segregation) between 90’-150’. Shortly after anaphase entry, cells completed the first cell cycle and split mother and daughter cells (new rise of unbudded category by 180’). We observed an initially similar cell cycle profile in the case of *top2-5* (Figure 1a, right panel), yet with an earlier G1-S transition [15]; S-phase entry at 30’-60’ and the peak of anaphase onset at 90’. As reported in our previous work, we seldom observed genuine H2A2-GFP anaphase bridges in *TOP2* (<10%) and they were transient in *top2-5* (∼60% of cells at 90’). Instead, binucleated cells were the major segregation phenotype by 120’, indicative of cytokinetic furrow ingression by that time point [15]. This phenotype may encompass either cells in anaphase/telophase or mother and daughter cells in the next G1 before completing physical separation [23]. Strikingly, and unlike *TOP2*, a clear uneven segregation of the H2A-GFP was evident in *top2-5* (Figure 1b, c). *TOP2* and *top2-5* also differed in that ∼20% *top2-5* mother cells rebudded without separation of the first mother and daughter (Figure 1c) [15,23].

**Figure 1.**
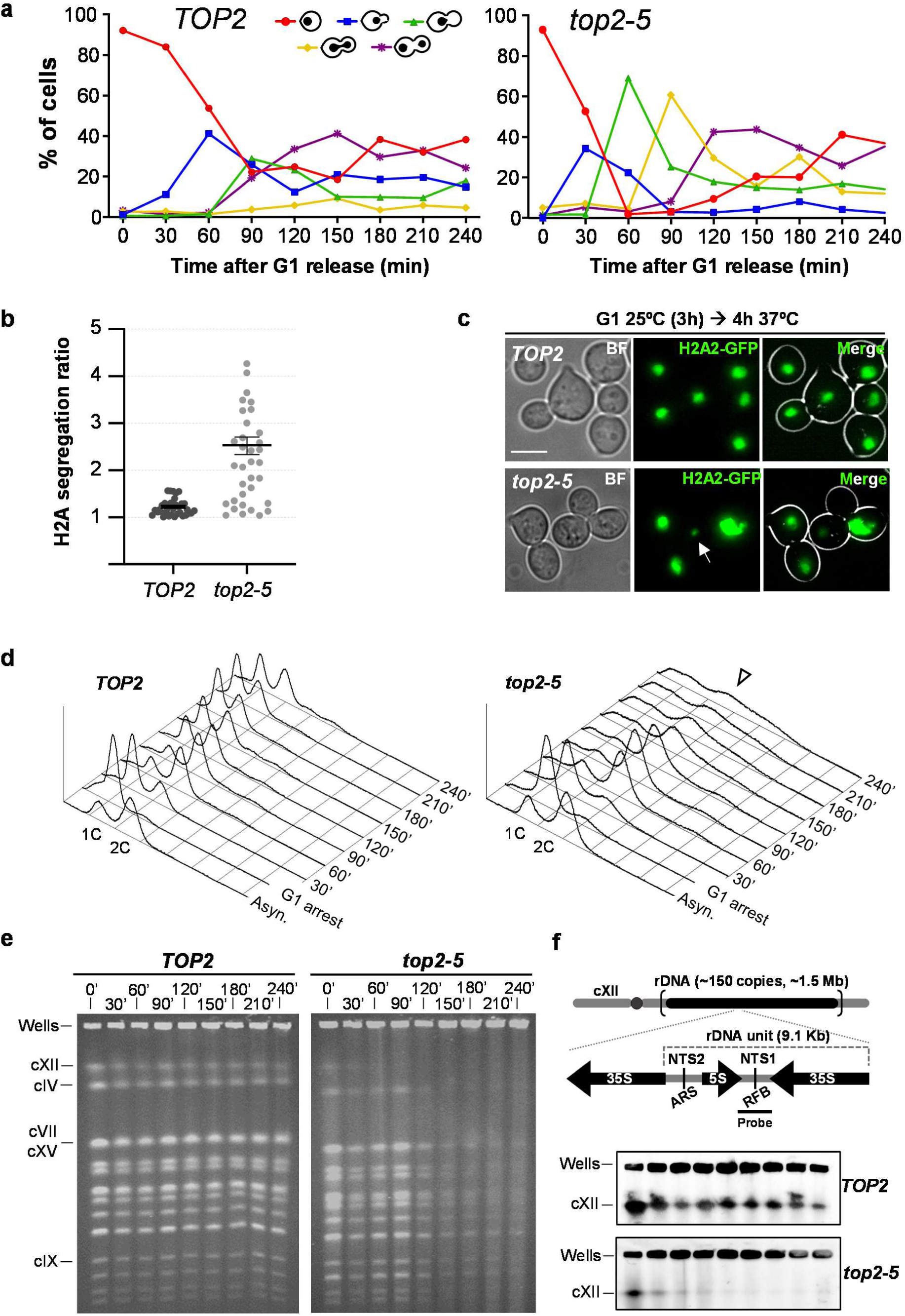
Chromosome integrity is compromised upon inactivation of Top2 with the *top2-5* thermosensitive allele. A synchronous G1 release experiment was performed for isogenic *TOP2* and *top2-5* strains. (**a**) Charts depicting the cell cycle progression under the microscope (both strains carry H2A2-GFP to label the nuclear masses). (**b**) Ratio of H2A2-GFP segregation among binucleated cells (n=34; 210’-240’ after the G1 release). The ratio is calculated by dividing the upper by the lower GFP signal in each pair of nuclei. (**c**) Micrographs of representative cells 4 h after the G1 release. The arrow points to a massive uneven segregation of the H2A2-GFP signal in *top2-5*. (**d**) Flow cytometry (FACS) analysis of the DNA content. Arrowhead highlights the flattened DNA content observed in *top2-5* at later time points. (**e**) Ethidium bromide (EtBr) staining of whole chromosomes resolved by Pulsed-Field Gel Electrophoresis (PFGE). Note how in the *top2-5* strain the cXII in-gel signal disappears shortly after the G1 release, whereas the other chromosome bands get fainter after 120’. (**f**) Southern blot profiles of the same PFGEs with a specific probe against the non-transcribed spacer region 1 (NTS1) of the ribosomal DNA array (rDNA) on the cXII (schematic on the top).

FACS analysis confirmed that both strains have a timely cell cycle, at least until anaphase onset (Figure 1d); S-phase entry (drop of 1C, rise of 2C DNA content) at 30’-60’. Cytokinesis and cell separation took place at 120’ in *TOP2* (new rise of 1C peak). Likewise, *top2-5* executed cytokinesis at 150’-180’ (drop of 2C peak); however, the outcome of such point-of-no-return was catastrophic. The 2C peak did not revert to the typical 1C DNA content but instead extended from <1C to >2C DNA content, indicating massive uneven segregation of the genetic material, as suggested by the H2A2-GFP microscopy.

When we performed the PFGE analysis, we observed that the chromosome staining pattern in *TOP2* was constant throughout the time course (Figure 1e, left). By contrast, there were two waves of decreased chromosome staining in *top2-5* (Figure 1e, right; Figure S1). In the first wave, the signal from largest chromosome, chromosome XII (cXII), decreased coincident with the cells transiting through S-phase. The second wave occurred after the nuclear division but before the decrease of the 2C peak by FACS (t=120’-240’), and was marked by a sharp and profound decrease of all chromosome staining (the in-gel signal dropped ∼90% in less than 1 h; Figure S1). We found that decrease in chromosome staining in *top2-5* persisted through different sample preparation conditions (Figure S2). Furthermore, the decrease in chromosome staining was not observed if cells were maintained in G1 through an equivalent incubation regime (6h from the induction of the G1 block) (Figure S3).

The lack of visible chromosomes in a PFGE can be due to three major causes: chromosome breakage, chromosome degradation and chromosomes with branched structures that keep them trapped into the loading well [7,24,25]. In order to distinguish between these possibilities, we did a Southern blot with a probe for the rDNA in cXII (Figure 1f & S4). We observed a strong cXII signal in the wells for the *top2-5* mutant. No signs of broken cXII (fast-migrating smear, e.g.) were noted (Figure S4). These results suggest that cXII gets trapped in the loading well. We did not observe signs of DNA degradation over the 4h time course without Top2, even after the massive missegregation of the genetic material (Figure S5).

We previously compared the *top2-5* allele with the broadly-used *top2-4* allele [15]. We found that *top2-5* transits through anaphase faster and more synchronously than *top2-4*, so that *top2-5* appears to be a better allele for cell cycle studies at this late stage. However, in order to address if our observations were specific to the *top2-5* allele, we checked the PFGE pattern in an isogenic *top2-4* strain. We found the same steady disappearance of all yeast chromosomes, yet to a lesser extent (Figure S6).

Overall, we conclude that Top2 inactivation brings about an unreported shift in the behaviour of all yeast chromosomes in a PFGE. In a synchronous cell cycle, this shift takes place in late anaphase, near or after cytokinesis.

### Cytokinesis is not responsible for the *top2-5* structural chromosome change revealed by PFGE

The general loss of chromosome bands in the *top2-5* PFGE from 120’ onwards could simply be due to cells completing a devasting cytokinesis, as both microscopy and FACS strongly suggest. Cytokinesis would break chromosomes at the anaphase bridges; and these broken chromosomes could get trapped in the PFGE well during DSB end resection [25]. Of note, we and others have shown that cytokinesis is a point of no return during Top2 inactivation [15,17]. In order to test this hypothesis, we used a *top2-5 cdc15-2* double ts mutant that blocks cytokinesis and cell progression beyond telophase because of the lack of the Mitotic Exit Network (MEN) kinase Cdc15 [2,15]. We repeated the time course and analysed samples by fluorescence microscopy, FACS and PFGE, comparing *top2-5 cdc15-2* with its *TOP2 cdc15-2* counterpart (Figure 2). As expected, cells from both strains arrested in telophase, as can be seen by microscopy (dumbbells prevailed) and flow cytometry (2C peak prevailed) (Figures 2a, b & c). As previously reported [15], *top2-5 cdc15-2* formed histone-labelled anaphase bridges (up to 90% of cells by 210’), whereas two equally segregated histone masses was the major outcome in the *TOP2 cdc15-2* (Figures 2a & b). Strikingly though, the PFGE of the *top2-5 cdc15-2* showed the same decrease intensity of the chromosome bands (Figure 2d), as we observed in *top2-5 CDC15*. Specifically, (i) in-gel cXII quickly disappeared (60’-90’), and (ii) all other chromosomes bands decreased at 120’-150’. We therefore conclude that the structural chromosome changes revealed by PFGE are not simply a consequence of the breakage of *top2-5* anaphase bridges by cytokinesis.

**Figure 2.**
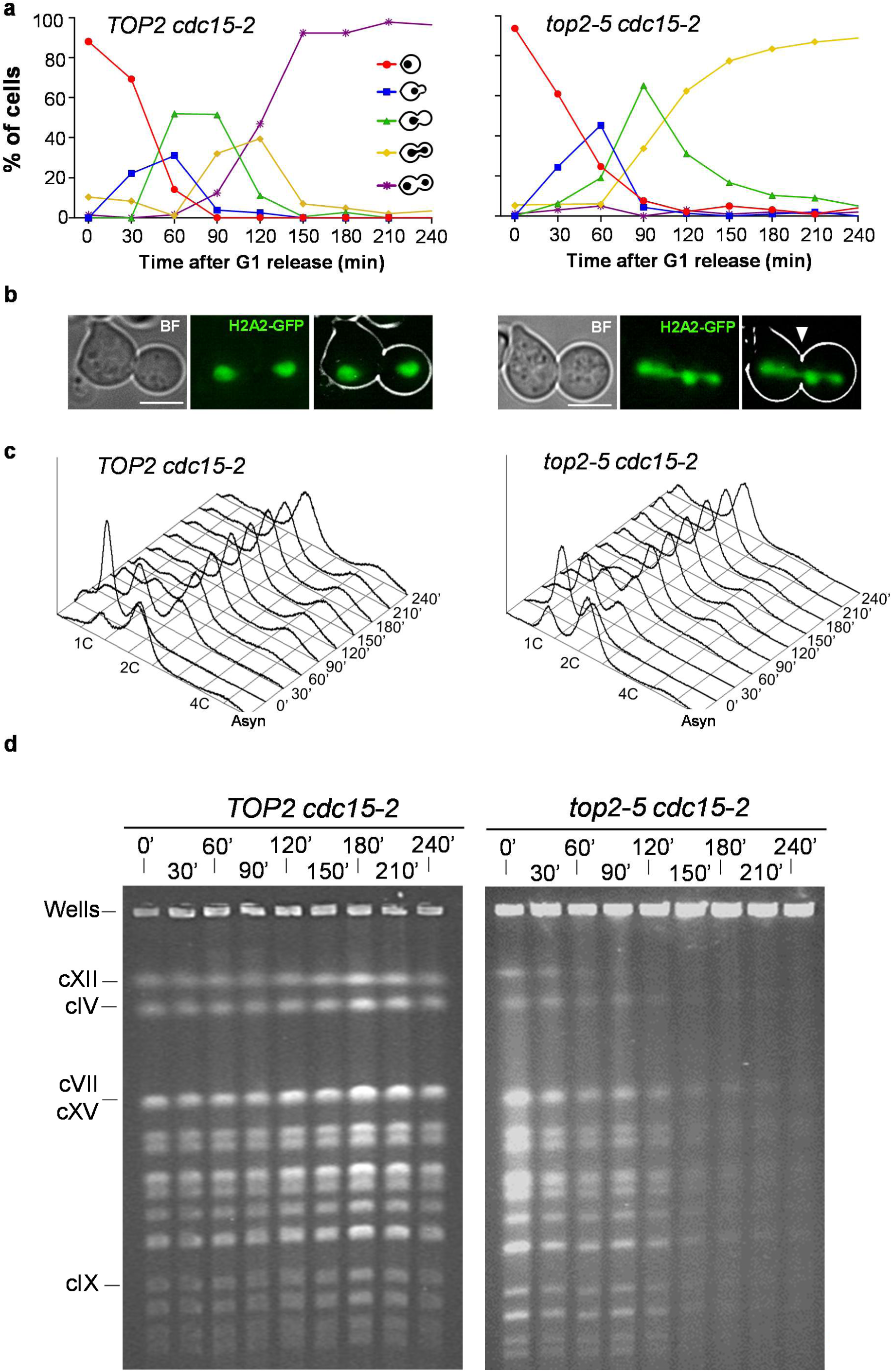
The loss of chromosome integrity in *top2-5* is not a consequence of the mitotic catastrophe that happens upon cytokinesis. A synchronous G1 release experiment was performed for isogenic *TOP2 cdc15-*2 (left panels) and *top2-5 cdc15-*2 (right panels) strains. (**a**) Charts depicting the cell cycle progression under the microscope. (**b**) Micrographs of representative cells 4 h after the G1 release. The arrowhead points to the characteristic massive H2A2-GFP anaphase bridge observed in *top2-5 cdc15-2*. (**c**) FACS analysis of the DNA content. (**d**) EtBr staining of whole chromosomes resolved by PFGE. Note how chromosome behaviour of the *top2-5 cdc15-2* strain resembled that of *top2-5*.

### Assessment of unfinished replication in *top2-5*

Since cytokinesis is not required for the loss of chromosome bands in PFGE, we tested whether long-lasting DNA-DNA sister chromatid junctions are responsible [7,24,25]. The timing of the 2C peak by FACS suggests that there is no a major delay in ongoing replication; however, certain late replication intermediates in *top2-5* might change the chromosome structure in such a way that cannot enter the PFGE. Indeed, previous results showed that yeast cells deficient in Top2 struggle to complete replication and accumulate late replication intermediates at replication termini sites [18]. Thus, we first tested whether *top2-5* accumulates late replication intermediates by performing neutral-neutral two-dimensional electrophoresis (NN-2D). This technique can detect the presence of branched DNA structures and classify them into replication-like (Y-shaped) and recombination-like (X-shaped) intermediates [27]. We began studying the rDNA since replication terminates at the well-defined RFBs and, indeed, we found enrichment of Y shapes in *top2-5 cdc15-2* (Figure 3a). The fixed Y-shaped structure at the RFB (spot “a” in Figure 3a) and other longer Ys that bypass the RFB into the adjacent 3’ end of the 35S gene (“b” in Figure 3a) were particularly enriched. In addition, other spots onto the X-shaped structures also accumulated (“c” and “d” at the spike in Figure 3a), which might correspond to two very close converging Ys as they cannot branch-migrate along the spike.

**Figure 3.**
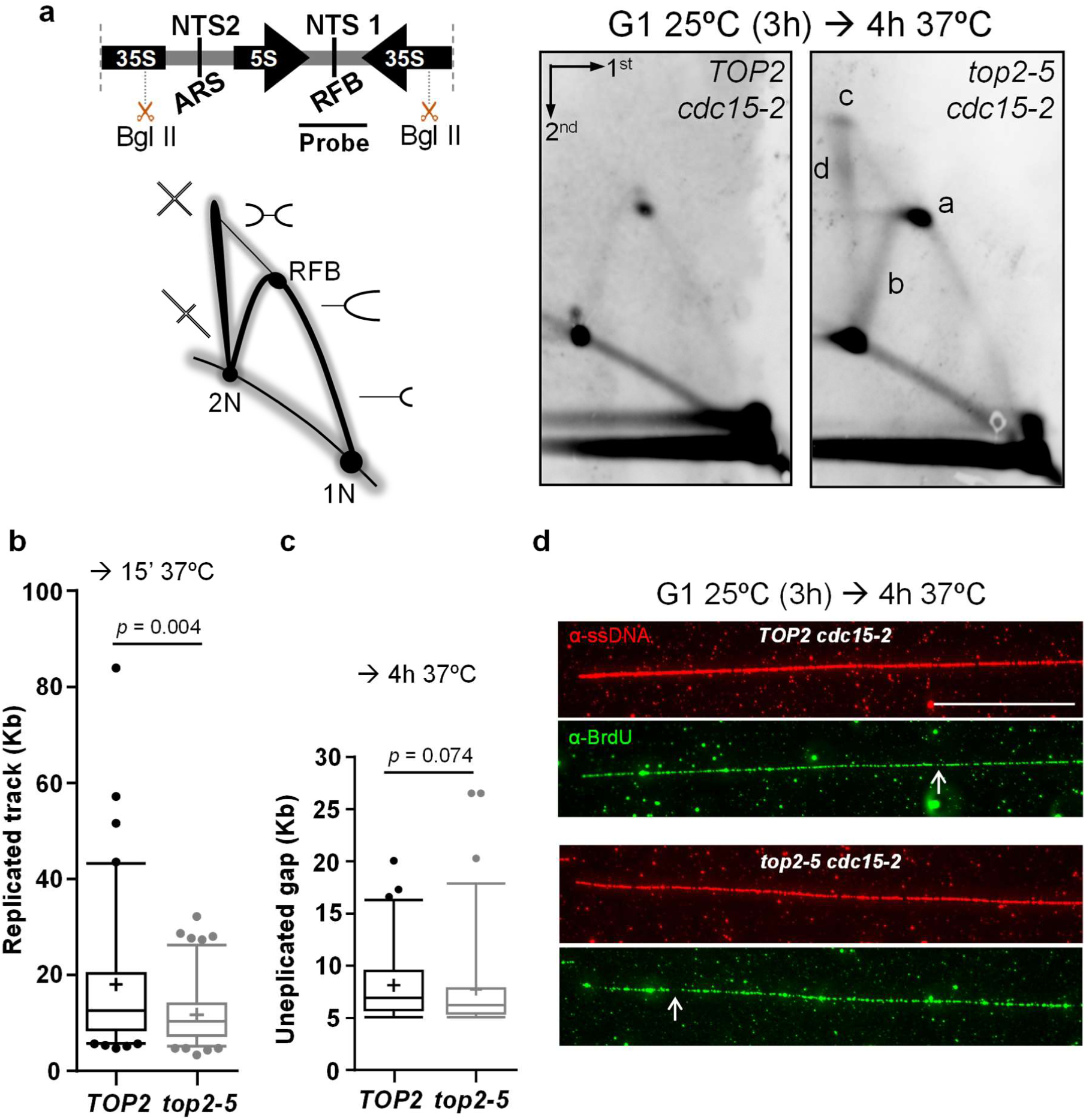
Assessment of the presence of underreplicated chromosomes in *top2-5*. Synchronous G1 release experiment were performed for isogenic *TOP2 cdc15-*2 and and *top2-5 cdc15-*2 strains. (**a**) The presence of persistent replication and recombination-like intermediates were evaluated by neutral, neutral two-dimensional electrohpresis (NN-2D) for the NTS1 region of the 9.1 Kbps unit of the rDNA. On the top left, schematic of the rDNA unit with its main features and relative position of the BglII sites and the probe; a representation of the theoretical branched forms visualized by NN-2D is depicted underneath. NTS1 & 2, non-transcribed regions 1 & 2, respectively; ARS, autonomous replicating sequence; RFB, replication fork block. On the right, NN-2D of both strains 4h after the G1 release into 37 °C. Note that four structures were enriched in *top2-5 cdc15-2* relative to *TOP2 cdc15-2*: the Y-shaped intermediate stalled at the RFB (“a”), the long Y-shaped intermediates that pass such block (“b”), the intermediate with highest mass and symmetry (“c” spot, probably a double Y), and the X-shaped spike (“d”). (**b**) Combing analysis of the beginning of replication with or without Top2 activity (15’ after the G1 release). (**c**) Combing analysis of the completion of replication for DNA fibers obtained 4 h after the G1 release. (**d**) Representative pictures of the 4h immunofluorescence against DNA (red) and incorporated BrdU (green). White arrows point to the small greenless gaps observed in fibers coming from both strains. The scale bar represents 50 Kbps.

For chromosomes other than XII, it is difficult to determine where two replication forks converge and is expected to be variable in the cell population. Thus, we opted for an alternative technical approach based on measuring underreplicated gaps on extended DNA fibers. These gaps would suggest the presence of two converging RFs (double Ys) in a single-molecule analysis. For that purpose, we transferred the *top2-5* and *cdc15-2* alleles to a strain suitable for the DNA combing technique [28]. Although we observed a slower replication fork progression at the beginning of *top2-5* S-phase (Figure 3b), we could not see differences between the *top2-5 cdc15-2* and *TOP2 cdc15-2* strain in late anaphase (Figure 3c & d). Fibers appeared almost fully replicated in both cases; i.e., green (replicated DNA) and red (DNA) signals extensively overlapped. Although there were some gaps in the BrdU signal on the DNA fibers, there were no differences between the strains in terms of underreplicated percentage (5.85% and 5.34% for *TOP2* and *top2-5*, respectively) and track length (Figure 3c). The number of gaps (greater than5 Kbs) was also equivalent in the two strains (72 & 69 gaps in 10 Mbs for *TOP2* and *top2-5*, respectively). The data suggest that unreplicated regions larger than 5 kbp do not accumulate in *top2-5*, although Y-structures at the rDNA RFB and X-structures at the rDNA do accumulate.

### Assessment of unresolved recombination intermediates in *top2-5*

The X-shaped molecules observed by NN-2D in the *top2-5* rDNA suggest that recombination intermediates might also contribute to the trapping of chromosomes in the PFGE well. HR is known to bypass stalled RFs and might help in completing replication in *top2* mutants [29]. In order to assess the contribution of recombination intermediates to the *top2-5* PFGE and NN-2D profiles, we used a genetic approach since such intermediates depend on the HR gene *RAD52* [30]. The triple mutant *top2-5 cdc15-2 rad52*Δ strain showed a similar cell cycle profile than *top2-5 cdc15-2*, including anaphase bridges as the end-point phenotype in telophase (Figure 4a) and similar replication timing by FACS (Figure 4b). Interestingly, the early loss of the cXII band was partly alleviated in this triple mutant (Figure 4c). However, the abrogation of Rad52 neither restored the late loss of chromosome bands in the PFGE nor affected the presence of X-shaped intermediates at the rDNA (Figure 4d). We conclude that recombination intermediates are unlikely to contribute to the *top2-5* anaphase decrease in chromosome bands by PFGE or to the X-shaped structures at the rDNA..

**Figure 4.**
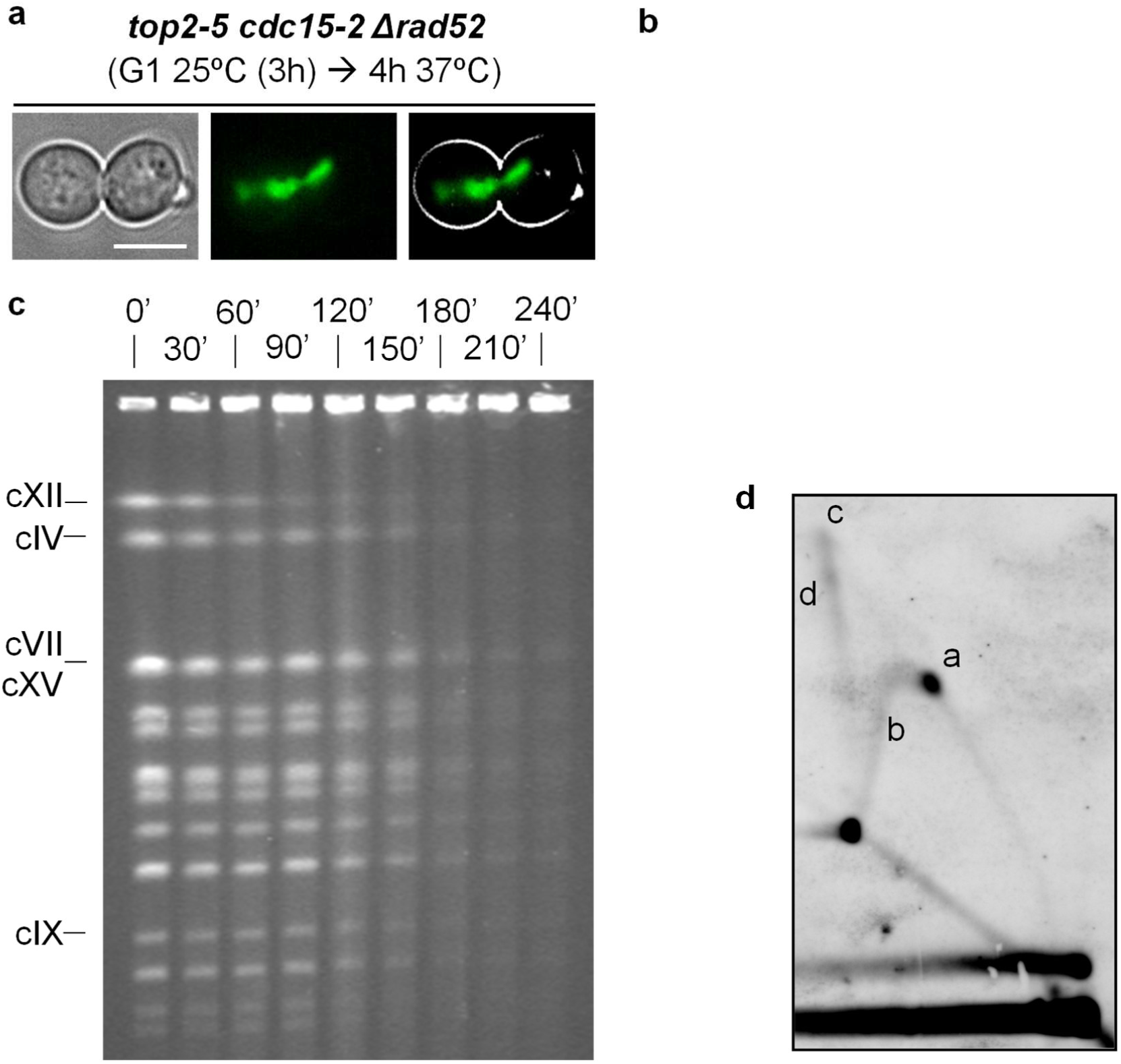
Chromosome integrity loss in *top2-5* is independent of the homologous recombination pathway. A synchronous G1 release experiment was performed for the *top2-5 cdc15-*2 Δ*rad52* strain. (**a**) Micrographs of a representative cell 4 h after the G1 release. Note it resembles *top2-5 cdc15-*2 with its characteristic H2A2-GFP anaphase bridge. (**b**) FACS analysis of the DNA content. (**c**) EtBr staining of whole chromosomes resolved by PFGE. Note how the late chromosome shift behaviour resembled that of the *top2-5 cdc15-2* strain. (**d**) NN-2D of the NTS1 region of the rDNA 4 h after the G1 release.

### The *top2-5* structural chromosome change revealed by PFGE is independent of *CDC14*-mediated processes in early anaphase

Since cytokinesis was not responsible for the loss of PFGE chromosome bands in *top2-5*, we wondered whether transition through anaphase played any role. Anaphase starts when all sister kinetochores are attached to opposite spindle pole bodies through the microtubule-based spindle apparatus [31]. At that point, cohesion between sister chromatids is lost and the spindle pulls sisters apart. Once cells enter anaphase, the master cell cycle phosphatase Cdc14 is activated through the FEAR network to promote spindle elongation [32,33], as well as resolution and condensation of the rDNA [34–37]. In addition, Cdc14 targets to the nucleus and activates the structure-specific endonuclease Yen1 [38–40], which can recognize and cut both Y-shaped and X-shaped molecules [38]. We reasoned that any of these Cdc14-mediated events could be responsible for the loss of chromosome bands observed ∼120’ after the G1 release. However, the *top2-5 cdc14-1* strain was indistinguishable from *top2-5 cdc15-2*, ruling out this possibility (Figure S7).

### The *top2-5* structural chromosome change revealed by PFGE does not depend on anaphase onset or on spindle force

Neither cytokinesis nor Cdc14-controlled anaphase events were responsible for the PFGE phenotype, yet it clearly takes place in anaphase as determined by comparing microscopy and PFGE (Figures 1, 2 and S7). We next checked whether the force of the mitotic spindle might trigger the loss of chromosome bands. Thus, we added nocodazole (Nz) to depolymerize the microtubules before *top2-5* cells reached anaphase. Noteworthy, Nz also elicits the activation of the spindle assembly checkpoint [41], leading to a cell cycle block at metaphase. The FACS pattern we observed, with the corresponding long lasting 2C peak in the single *top2-5* mutant (Figure 5a, upper FACS profile), confirmed this arrest. The presence of >95% of mononucleated dumbbell cells under the microscope (Figure 5b, lower left) further confirmed the Nz arrest. Strikingly, even though Nz prevented both anaphase onset and spindle pulling forces, the loss of chromosome bands still occurred, although with a 1 h delay compared to the previous examples (180’ versus ∼120’) (Figure 5c lanes 1-9).

**Figure 5.**
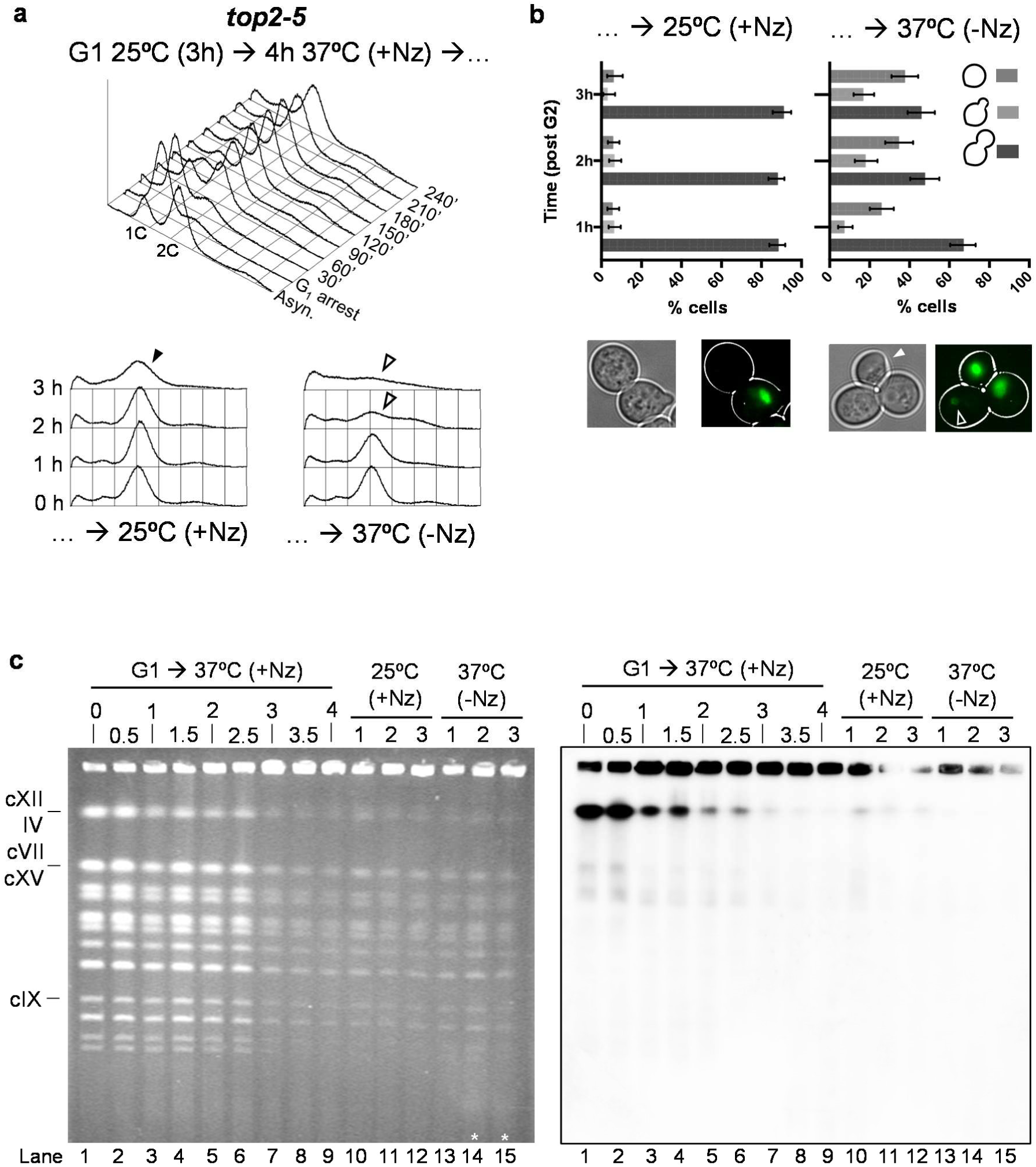
The late change in chromosome structure caused by the absence of Top2 is not a consequence of anaphase entry or mitotic spindle forces, is not reversed by Top2, and does not trigger a G2/M checkpoint. A synchronous G1 release experiment was performed for the *top2-5* strain under restrictive conditions for Top2 activity (37 °C). At the time of the G1 release, Nocodazole (Nz) was added to depolymerize the spindle and block cells in G2/M. After 4 h in Nz, the culture was split in two and incubated for another 3 h. One of them was shift back to 25 °C (i.e., re-activate Top2) while keeping the Nz block, whereas the other one was kept at 37 °C while removing Nz. (**a**) FACS analysis of the DNA content during the first 4 h of the G1-to-Nz cell cycle (on the top) and during the extra 3 h after the culture splitting (underneath). (**b**) Cell morphology analysis during the extra 3 h incubations. Below the charts, representative micrographs of each subculture after 3 h; i.e., a mononucleated dumbbell versus a dumbbell with an uneven segregation (hollow arrowhead) plus a postcytokinetic G1-like cell (white arrowhead). (**c**) EtBr-stained PFGE of the whole experiment (left) and the corresponding Southern blot against the rDNA (right). Note how (i) the late PFGE general shift still occurs in Nz, (ii) how reactivation of Top2-5 does not reshape the chromosomes to enter the gel, and (iii) how the shift does not trigger a G2/M, so that a mitotic catastrophe still takes place after Nz removal (asterisks in lanes 14 & 15).

### The *top2-5* structural chromosome change revealed by PFGE cannot be reverted by re-activation of Top2

Having found that the loss of chromosome bands can also happen in metaphase-arrested cells after prolonged absence of Top2, we next checked whether re-activation of Top2 could restore the chromosome bands. For that purpose, we shifted the temperature down to 25 °C while maintaining the cells in Nz. In this manner, we avoided any downstream events in anaphase that might influence the experimental outcome. Indeed, cells remained in metaphase after the temperature shift for at least 3 h (Figure 5a, lower left FACS profile; and 5b, left chart). Surprisingly, Top2 re-activation did not restore the structural integrity of the chromosomes (Figure 5c, lanes 10-12). We conclude that upon prolonged Top2 depletion, chromosomes change their structure irreversibly.

### G_2_/M checkpoints are blind to the *top2-5* structural chromosome change

The persistent loss of chromosome bands in Nz-arrested cells allowed us to assess whether the structural chromosome change triggers a G_2_/M checkpoint. This question could not be answered in cycling cells as the change happens in anaphase. However, once we observed the loss of chromosome bands in Nz, if a G_2_/M checkpoint was triggered we would see a delay in anaphase onset after Nz removal. However, when Nz was removed we observed not only anaphase progression but also cytokinesis and the corresponding mitotic catastrophe (Figure 5a, lower right FACS profile; and 5b, right chart).

In addition, we ruled out that the absence of a checkpoint response after the *top2-5* PFGE shift was a consequence of Top2 itself being a sensor/mediator of such checkpoint(s). Both hydroxyurea (HU) and methyl methanesulfonate (MMS) still arrested the *top2-5* strain in G_2_/M in restrictive conditions (Figure S8). HU creates replicative stress through formation of ssDNA behind the RFs, whereas MMS leads to RF stalling in addition to ssDNA gaps behind the RFs [5].

### A GFP-based candidate screen of DNA damage and checkpoint proteins further indicates that *top2-5* cells do not detect the structural chromosome change as chromosomal damage

In addition to the previous set of experiments, we conducted a GFP-based screen of DNA damage proteins known to either form foci or increase their nuclear content upon DNA damage [42]. This screen was undertaken at the *top2-5 cdc15-2* block and included the Dpb11-yEmRFP as an additional reporter. Dpb11 is known to get enriched at certain types of anaphase bridges [43]. As a control, we included a *TOP2 cdc15-2* strain also blocked in telophase. We observed an increase in nuclear foci in the *top2-5 cdc15-2* block for the DNA replication stress markers Lcd1/Ddc2, Rfa2 and Rfa3, the latter two belonging to the ssDNA binding RPA complex (Table 1) (Figure S9 and S10). The foci were predominantly present along the anaphase bridge. However, there were no differences for other important replication stress reporters such as Ddc1, Dpb11, Dna2 and Rad5. Importantly, there was no increase in foci of proteins involved in HR such as Rad51, Rad52 and Rad54. Altogether, we conclude that more ssDNA is present in cells blocked in telophase after passing through a cell cycle without Top2. However, this higher ssDNA level does not elicit an efficient DDR. Incidentally, there were RPA and Dna2 foci in *TOP2 cdc15-2* cells. Nearly 50% of telophase-blocked *TOP2 cdc15-2* cells had one Rfa2/3 focus.

**Table 1.**
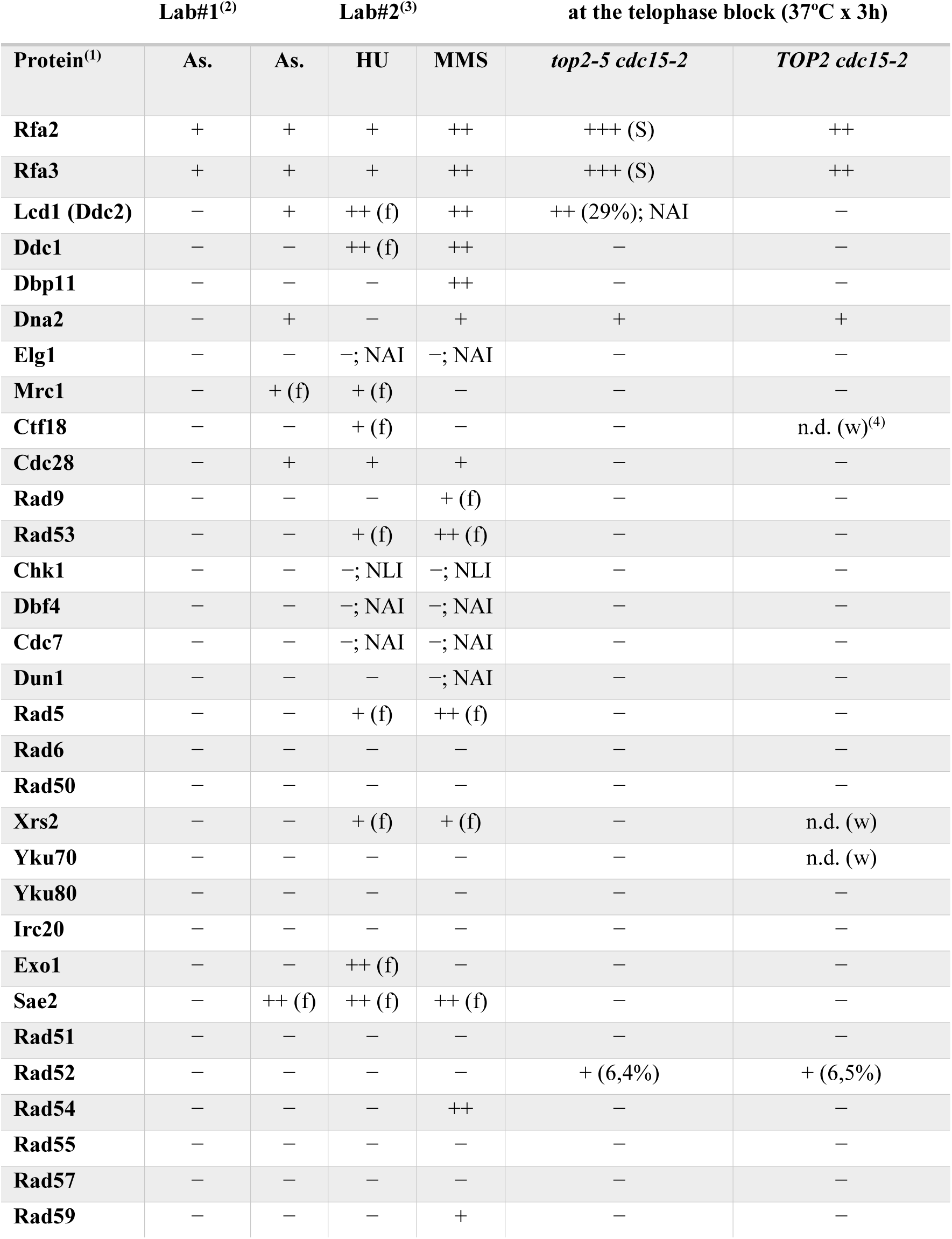
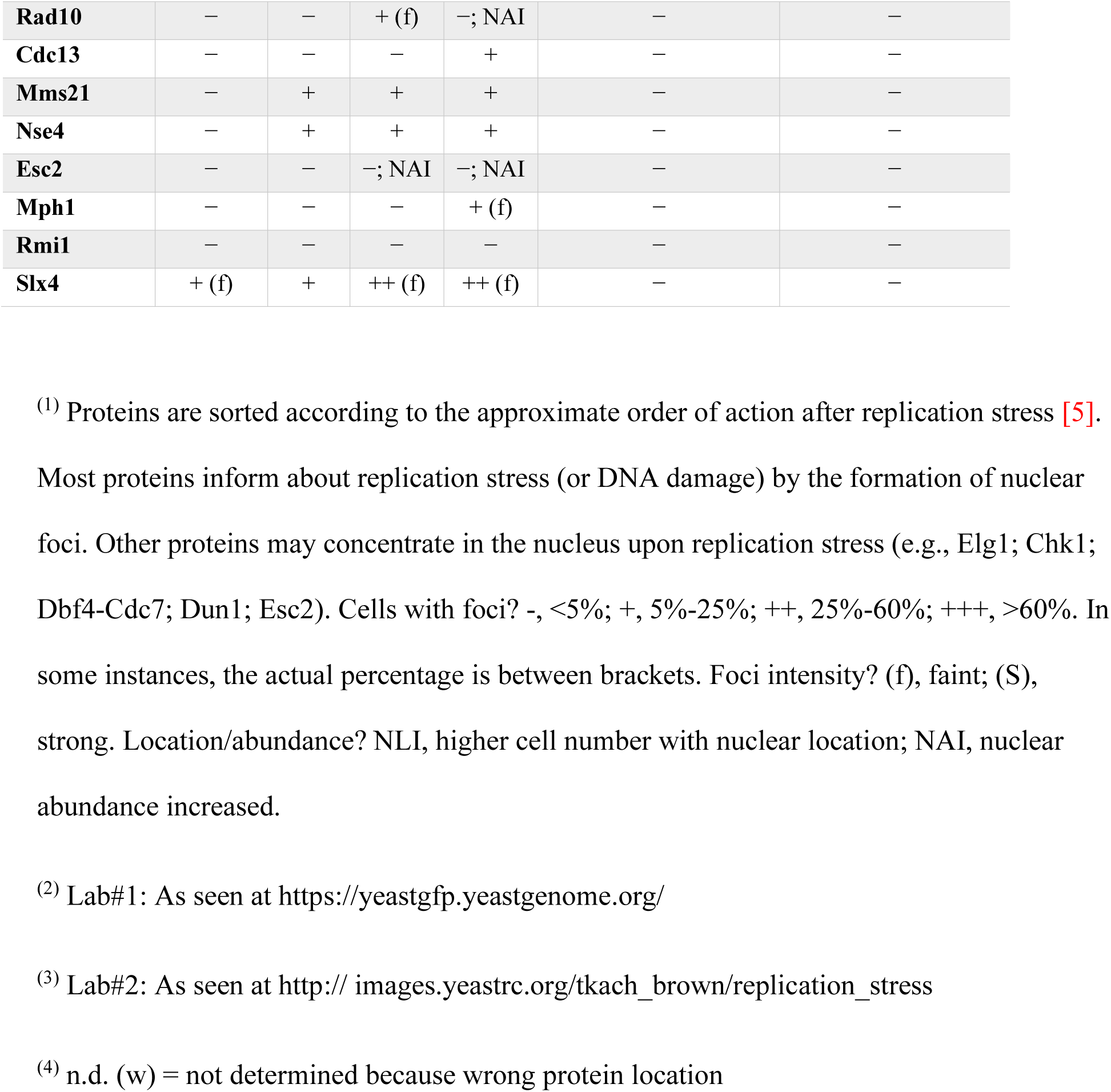
Location pattern in telophase-blocked cells of DNA damage checkpoint and repair proteins after going through a synchronous cell cycle with or without Top2.

### In vitro treatment with T7 endonuclease partly releases *top2-5* chromosomes from the PFGE well

Previous studies on the well-bound DNA after PFGE showed that it often carries DNA molecules which are larger than expected, and that they form unusual multifiber structures such as starbursts and comets [44]. Many of these studies were undertaken with simple circular bacterial or mitochondrial DNA (mtDNA), as short as 20 Kbs, and the PFGE well trapping phenomenon has been attributed to rolling circle replication (5’ flap branched structures), abundance of ssDNA, and origami-like branched supermegabase organization [45,46]. The apparent increase in ssDNA in the *top2-5 cdc15-2*, as well as the branched structures observed at the rDNA, led us to test how enzymatic treatment of the DNA plugs might alter the subsequent chromosome migration in a PFGE. Samples from 0’ (G_1_ block) and 180’ (telophase block) were treated with four enzymes that modify ssDNA and/or branched DNA (Figure 6). Firstly, Klenow fragment of *Escherichia coli* DNA polymerase I can fill ssDNA gaps while creating 5’ flaps (a type of branched DNA) by displacing the downstream dsDNA once the gap is filled. Secondly, S1 and Mung bean endonucleases cleave DNA at ssDNA gaps. Finally, T7 endonuclease I cleaves branched DNA and heteroduplex dsDNA.

**Figure 6.**
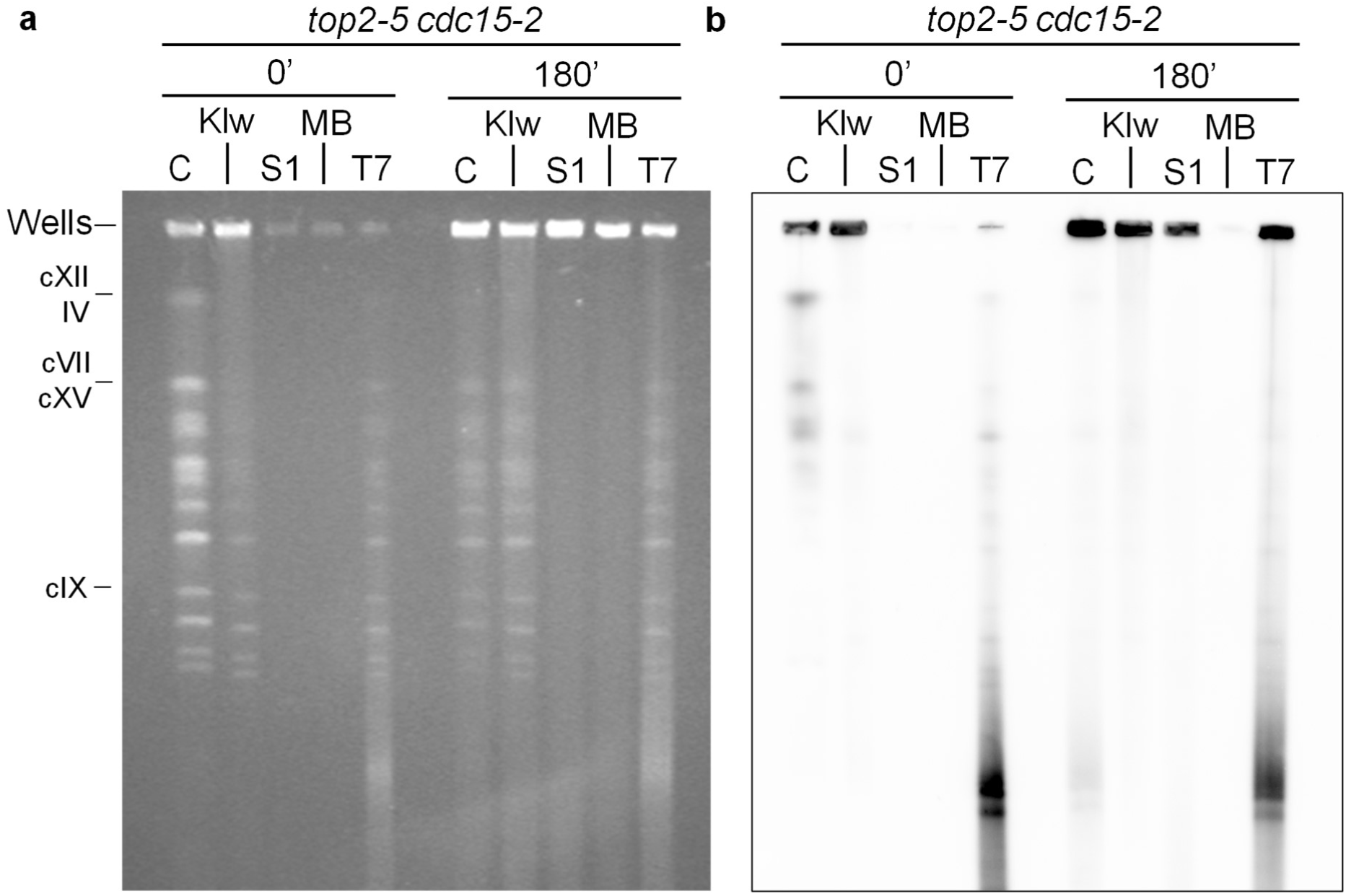
In vitro treatment with the structure-selective T7 endonuclease I partly suppresses PFGE chromosome entrapment. A synchronous G1 release experiment was performed for isogenic *TOP2 cdc15-*2 and *top2-5 cdc15-*2 strains. Plugs from 0’ (G1 block) and 180’ after the release at 37 °C were treated as indicated before running a PFGE. (a) EtBr-stained gel. (b) Southern blot against the rDNA. Note that Klenow treatment entraps G1-blocked chromosomes while T7 endonuclease releases entrapped rDNA from the 180’.

Treatment of plugs with S1 and Mung Bean extensively degraded DNA from both plugs. This may suggest that ssDNA gaps are already present at the G1 block. However, it is more likely that the DNA in the plugs contains nicks, either natural or acquired during extraction. Nicked DNA (nDNA) is a known substrate of both nucleases and generate DSBs by a counternick mechanism [47]. In either case, ssDNA and/or nDNA did not preclude G_1_ chromosomes from entering the PFGE. Treatment with Klenow led to entrapment of the G_1_ plug but did not modify the degree of retention of the 180’ sample (Figure 6a). The entrapment of the G_1_ sample is consistent with the possibility of Klenow creating branched structures through strand displacement from ssDNA or nDNA. Lastly, T7 endonuclease trimmed most chromosomes into a fast migrating smear (Figure 6a). This happened in the G_1_ sample but, importantly, it also occurred in the 180’ plug up to a level where both samples had equivalent DNA entering the PFGE. In other words, T7 endonuclease releases the DNA entrapped in the well in the *top2-5 cdc15-2* telophase block. The DNA release from the well included the rDNA as seen by Southern blot analysis (Figure 6b). The results obtained with Klenow and T7 strengthen the argument that branched structures account for the retention of chromosomes after prolonged absence of Top2.

## DISCUSSION

Top2 is the critical enzyme that removes sister chromatid intertwines prior to chromosome segregation in anaphase. The absence of Top2 is broadly recognized to lead to a type of mitotic catastrophe whereby anaphase bridges, which arise from these intertwined chromatids, are severed during cytokinesis. Altogether, the results shown in this paper confirmed that *top2-5* also leads to a mitotic catastrophe; we here demonstrate this by microscopy, complementing our previous work [15,23], but also by FACS and by PFGE. The mitotic catastrophe in *top2-5* seems greater than other previously studied *top2-ts* alleles, at least by FACS analysis and microscopy [15,17,48], and might be a feature of either the *top2-5* allele itself or the genetic background it is in [15]. However, the most shocking *top2-5* phenotype we introduce here is the disappearance of chromosome bands in a PFGE at a stage prior to the mitotic catastrophe. The overall DNA content we quantified by FACS, the absence of in-gel smears and broken rDNA in the Southern blots, and the re-appearance of in-gel signals after in vitro treatment with T7 endonuclease, strongly suggests that the disappearance of bands correlates to entrapment of chromosomes in the loading well (Figures 1, 2, 6, S4-S6). Moreover, DNA release from the plug was also accomplished by digestion with BglII for the NN-2D analysis (Figures 3 and 4). We distinguished two classes of chromosomes affected by the PFGE retention: the rDNA-bearing chromosome XII and all the other 15 chromosomes. The former persistently gets well-bound since the beginning of the S phase, which is compatible with, at least, unfinished replication (protracted Y-shaped DNA at the RFB in *top2-5 cdc15-2*; Figure 3 and 4). By contrast, chromosomes other than XII manifest the same PFGE behaviour much later in the cell cycle; after replication appears to have ended by both FACS and PFGE (Figures 1, 2, 4, 5, S1, S3-S7). A closer look at this late PFGE behaviour suggests it is independent of transition between cell cycle stages (i.e., S-phase to metaphase to anaphase) and the key spatial and molecular changes that occur in such transitions (e.g., activation of structure-specific endonucleases, spindle pulling forces, hypercondensation by Cdc14, etc) [2,49]. Rather, it seems there is a fixed time window between the end of S-phase (as determined by FACS) and the PFGE shift (∼1-2 h). Thus, in synchronous cell cycle cultures the loss of chromosome bands takes place well within anaphase (Figures 1, 2 & S7), and the loss of chromosome bands is present in Nz-treated cells as well (Figure 5). Importantly, this latter experiment led us to conclude two important features of the loss of chromosome bands in PFGE; (i) no G_2_/M checkpoint is activated after the loss of chromosome bands; and (ii) the loss of chromosome bands is irreversible with respect to Top2 activity.

There are four known causes of chromosomes entrapment in the loading well during PFGE: linear chromosomes larger than 10 Mbs [50], relaxed circular chromosomes larger than 100 Kbs [51], the presence of chromosomes with branched structures [7,24,25,52], and the presence of large portions of ssDNA [45,46]. Branched structures and ssDNA physiologically arise during replication and DNA repair through HR, and we believe they are the underlying cause of the observed loss of chromosome bands. Whereas a theoretical and experimental framework exists to explain the relationship between PFGE trapping and chromosome size [53], the causes of why chromosomes carrying DNA branches or ssDNA gaps do not enter PFGE remain undetermined. Because of this lack of knowledge on the PFGE technique, we cannot fully draw at present the postreplicative pathway to whatever structures preclude the affected chromosomes from entering the PFGE. From results we present in this work (Figure 3, 6, S9 and S10; Table 1), it appears that both ssDNA and replicative branched structures are already present in chromosomes that, nonetheless, enter a PFGE. For instance, gaps of unreplicated DNA were observed by combing in the *TOP2 cdc15-2* block (Figure 3). Although this may reflect a limitation of the combing technique for the purpose of quantifying underreplication, it is remarkable that a very recent report shows that chromosomes are not fully replicated in a *TOP2 cdc15-2* block [54]. Likewise, we have shown that sister chromatids are somehow connected with each other in the *TOP2 cdc15-2* block and form retrograde anaphase bridges after DSBs [55]. In addition, cytological markers of ssDNA and replication stress are present in the *TOP2 cdc15-2* block, although there is an increase in the *top2-5 cdc15-2* block (Table 1; Figures S9 and S10). One of this markers, the RPA complex, has been recently seen by others in the *TOP2 cdc15-2* block [54]. Because we could not see a difference in the amount of underreplicated material between *TOP2 cdc15-2* and *top2-5 cdc15-2* (Figure 3), we speculate that it is a late modification of the remaining Y-shaped branches in the absence of Top2 what triggers the PFGE shift. This modification is not sensed as DNA damage (Figure 1, 2 and S8) and does not depend on Rad52-driven HR (Figure 4). An interesting possibility is the eventual regression of converging RFs into four-way HJ-like chicken foot structures (Figure 7a), which are particularly enriched when cells cannot sense RF problems [56]. This scenario is compatible with the presence of Rad52-independent NN-2D X-shaped signals (Figure 4). Moreover, RF regression is expected for *top2* mutants as positive supercoiling accumulates ahead of the converging RFs at replication termini [57,58]. Where are these persistent RFs? We have shown that the RFB at the rDNA locus is one of these (Figure 3 and 4), but this only accounts for chromosome XII. We envisage that other difficult-to-replicate regions shared by all chromosomes may be involved. These regions would include centromeres, G-quadruplex, fragile sites, transposable elements, non-coding RNAs and subtelomeric regions [18,54,59–61]. Very recently, another paper has shown that pericentromeric regions accumulate DNA damage markers during S-phase [62]. Again, this damage does not appear to arrest the cell cycle [63]. A corollary of these assertions is that the traditional claim for ongoing replication as one cause of chromosome entrapment during PFGE should be revised. Perhaps, chromosomes get entrapped not by having RFs but by ensuing RF modifications.

**Figure 7.**
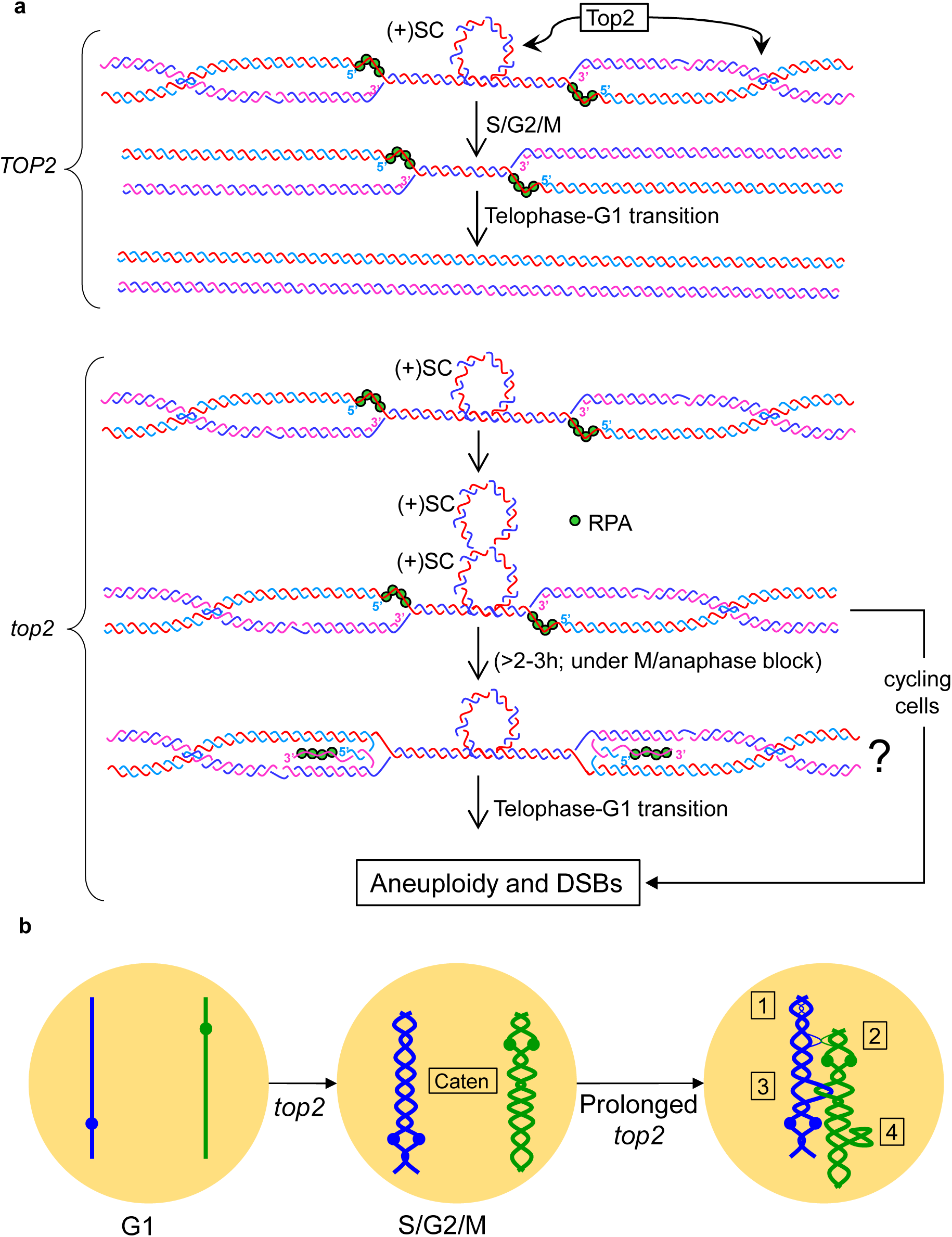
Models of Top2-irresolvable changes on the chromosome structure occurring upon prolonged Top2 deficiency. (**a**) When replication is ending and two converging forks are approaching, critical positive supercoiling arise ((+)SC) in between and precanenanes behind. Supercoiling interferes with replication completion, especially in difficult-to-replicate regions. Top2 deals with both topological problems, replication can finish, and sister chromatids are resolved for segregation. Part of these difficult-to-replicate regions finish replication during the telophase-G1 transition, according to a recent report. In the prolonged absence of Top2, difficult-to-replicate regions accumulate an excess of topological problems that shift the chromosome structure so that chromosomes get trapped in a PFGE. Based on previous literature and the findings shown herein, we speculate that an excess of (+)SC may turn into reversed forks, and that these structures, or derivatives thereof, lead to PFGE entrapment. (**b**) Sister chromatid catenations arise during replication. Without Top2, this class of intertwining persists. Meanwhile, other DNA metabolic processes go on, such as transcription and condensation, and these might exacerbate topological problems, so that other intertwining classes arise: 1, sister chromatid hemicatenanes; 2, chromosome hemicatenanes; 3, chromosome catenations; and 4, knots. Both models are not mutually exclusive. RPA, ssDNA-binding replication protein A complex.

Finally, we cannot rule other possibilities such as massive topological intertwinings, or even knots, between different chromosomes after prolonged Top2 absence (Figure 7b). Catenations between the replicated sister chromatids is the immediate consequence of the Top2 absence; however, within the constrained space of the nucleus, and with other important dynamic events taking place in G_2_/M/anaphase, such as transcription and condensation, it is difficult to envisage how cells avoid cumulative topological problems without Top2. Of note, previous studies with *top2-ts* showed vastly interlocked/knotted plasmids [16,17,64]. Massive interchromosomal intertwinings/knotting may form a chromosome web that trap chromosomes. Alternatively, cumulative topological stress could promote dsDNA unwinding towards ssDNA that, followed by ectopic or sister re-annealing, would create four-way DNA origamis. In addition, ssDNA tracks may be interlocked by type IA topoisomerases (Top3), forming hemicatenanes. These solutions are indeed compatible with the presence of more ssDNA in *top2-5 cdc15-2* while not observing larger underreplication gaps than the control *TOP2 cdc15-2*. In addition, it predicts Rad52-independent X-shaped molecules at the rDNA and lack of sensing by the DNA damage checkpoints. The presence of persistent late replication intermediates and complex topological intertwinings as explanations of the PFGE are not mutually exclusive. Indeed, knotting and hemicatenanes have been related to failure to finish replication [65,66].

In conclusion, the lack of Top2 postreplicatively modifies the structure of all yeast chromosomes in a way that diminish their ability to run in a PFGE. Future work on the actual physical nature of the DNA molecules trapped in the PFGE, together with genetic screenings of modifiers, should clarify this late chromosome structural change and perhaps assign a novel role for Top2 in the chromosome biology field.

## MATERIALS AND METHODS

### Yeast strains and experimental conditions

All the strains used in this work are listed in Table S1. All strains were grown overnight in air orbital incubators at 25 °C in YEPD. Time course experiments were performed as follows: asynchronous cultures were adjusted to OD_600_ = 0.3 and then synchronized in G1 at 25 °C for 3 h by adding 50 ng/ml of alpha-factor. The G1 release was induced by washing the cells twice and resuspending them in fresh media containing 0.1 mg/ml of pronase E. Then, they were incubated at 37°C for 3-4 h. In time course experiments, samples were taken every 30 minutes for microscopy, FACS and PFGE.

### Microscopy and flow cytometry (FACS)

H2A2-GFP was analyzed by wide-field fluorescence microscopy as reported before [15]. Briefly, series of z-focal plane images (15–20 planes, 0.15–0.3 mm depth) were collected on a Leica DMI6000, using a 63x/1.30 immersion objective and an ultrasensitive DFC 350 digital camera, and processed with the AF6000 software (Leica). Scale bars in micrographs depict 5 μm.

DNA content by flow cytometry analysis (FACS) was done as previously described using a BD FACScalibur equipment [67]. An asynchronous culture of each strain growing at permissive temperature was used to calibrate the 1C and 2C peaks before reading the samples.

### Pulsed-field gel electrophoresis (PFGE) and Neutral, neutral two-dimensional electrophoresis (NN-2D)

Yeast DNA for PFGE and NN-2D was extracted in low-melting agarose (LMA) plugs in conditions known to avoid branch migration [7]. Briefly, 4 OD_600_ equivalents were embedded into a 0.5% (w/v) LMA plug and then digested with zymolyase (2 units, day 1), RNaseA (10 µg/ml; day 1) and Proteinase K (1 mg/ml; day 2). All digestions were carried out at 37 °C (including the Proteinase K step). Yeast DNA for combing analysis was also extracted in LMA but in different conditions (see below). In case of subsequent treatment of the agarose plugs with DNA modifying enzymes, Proteinase K was first inactivated with 1 mg/ml Pefabloc® (11429868001, Roche). For all enzymatic treatments, a further overnight incubation with 250 units of the corresponding enzyme was added. The enzymes were BglII (R0144M, New England Biolabs), S1 nuclease (M5761, Promega), Mung Bean nuclease (M4311, Promega) and T7 endonuclease I (M0302S, New England Biolabs).

PFGEs were carried out using a CHEF DR-III system (Bio-Rad) in two conditions. Most PFGEs followed the standard protocol we have used before [7], with minor modifications: 1% agarose gel in 0.5 × TBE buffer and run at 14°C for 40 h at 5 V/cm with an initial switching time of 47 seconds, a final of 170 seconds, and an angle of 120°. The chromosome bands in the gels were visualized with EtBr staining. Chromosome XII was identified by Southern blot using a fluorescein-labelled probe (Sigma-Aldrich, #11585622910) against the NTS1 region within the rDNA [7,26]. Detection was performed by chemiluminescence (Vilber-Lourmat Fusion Solo S equipment) using an anti-fluorescein antibody coupled to alkaline phosphatase (Sigma-Aldrich, #11426338910) and using CDP-star (Sigma-Aldrich, #11759051001) as the substrate.

NN-2Ds were also performed from agarose plugs. These were digested with BglII, run in two dimensions, and transferred to positively charged nylon membranes for Southern analysis as described before [7,26].

### DNA combing and immunodetection

DNA combing experiments were performed as described before [68]. Firstly, *top2-5* and *cdc15-2* alleles were transferred by PCR methods to a strain suitable for BrdU DNA labelling (Table S1) [28]. The resulting strains were first arrested in G1 at 25 °C by incubating with 5 µg/mL alpha-factor (*BAR1* genotype) for 3h. Cells were then released from G1 by adding Pronase E (100 µg/mL final concentration) and incubated 4h at 37 °C. Half an hour before Pronase addition, BrdU was added to the cultures at 400 µg/mL final concentration. Samples were taken 15’ after the release, which corresponds to half S-phase [68], and at 4h. Samples were immediately treated with 0.1% w/v sodium azide and cooled down to 0 °C to freeze replication.

DNA for combing was extracted in LMA plugs. Relative to the plugs prepared for PFGE and NN-2Ds, these plugs used the following settings: 4 x 10^7^ cells (∼1 OD_600_ equivalents), 0.5 U Zymolyase (2 days of incubation), no RNAse, and 3 days incubation with Proteinase K at 50 °C.

For DNA combing, the plugs were incubated at RT for 30 min with 1 μL of YOYO-1 (Molecular Probes, Y3601) in 150 μL of TE50 buffer (10 mMTris-HCl pH 7.0, 50 mM EDTA). Then, they were washed 3 times in TE50 buffer, and incubated twice for 5 min in 2 mL MES buffer (7:3 v/v MES hydrate:MES sodium salt 50 mM pH 5.7). After that, the plugs were melted at 72 °C for 20 min. The solution was transferred to 42°C for 15 min and incubated overnight in 3 U of β-agarase I (New England Biolabs, M0392). Next, the solution was heated to 72 °C for 10 min, cooled to room temperature (RT) and poured into the reservoir of the combing device. The silanized combing coverslips were incubated into the solution for at least 5 minutes, and then pulled out at a constant speed of 710 μm/s. Finally, they were incubated for 90 min at 60 °C, and mounted on a glass slide.

For immunodetection, the slides were dehydrated by sequential 5 min incubations with 70%, 90%, and 100% v/v ethanol at RT. They were air dried and the DNA was denatured in 1M NaOH for 25 min at RT. Next, the slides were washed 5 times in PBS and incubated for 5 min in PBS-T (PBS plus 0.05% v/v Tween-20). After that, 21 μL of blocking buffer (PBS-T plus 10% w/v BSA) were added and dispersed on the coverslip and the slides were incubated in a humidity chamber for 30 min at 37 °C. Next, the coverslips were removed by dipping the slides into PBS-T, and 21 μL of anti-BrdU solution (1:40 dilution in blocking buffer of the rat anti-BrdU antibody [AbD Serotec, MCA2060]) were added, dispersed and incubated (1 h) as the previous step. The same steps were carried out after that with the anti-DNA solution (1:50 dilution of the anti-DNA antibody [Millipore, MAB3034]) and the secondary antibodies solution (1:75 dilutions of Alexa Fluor 488 goat anti-rat and Alexa Fluor 546 goat anti-mouse [Molecular probes, A11006 & A11030, respectively]), always washing with PBS-T in between. Finally, 10 μL of ProLong Gold antifade reagent (Molecular Probes, 36930) were added and the samples were covered with a fresh coverslip.

Only DNA fibers larger than 100 kbs and markedly labelled with BrdU were included in the analyses. A low number of fibers (1% in *TOP2 cdc15-2* and 9% in *top2-5 cdc15-2*) were wholly devoid of BrdU. We considered that these fibers came from cells that did not enter S-phase after the G_1_ release, as we previously showed that ∼10% of the *top2-5* progeny at 25 °C may be inviable [23].

### Mini-screening of fluorescent DNA stress markers

The GFP-fusion library (Invitrogen, catalog number: 95702) was used as the basis for the mini-screening [69]. To introduce both the *top2-5* and *cdc15-2* alleles into the library, a series of PCR-based transformations were carried out to obtain the strain OQR84. This strain was also designed to carry the anaphase bridge reporter Dpb11-yEmRFP. The strain OQR84 was crossed with preselected strains from the GFP-fusion library and then haploids were selected for two genotypes: *DPB11-yEmRFP gene-GFP TOP2 cdc15-2* and *DPB11-yEmRFP gene-GFP top2-5 cdc15-2* (Table S1).

To carry out the screen, cells were grown overnight in 96-well plates at permissive temperature (25°C) in synthetic complete medium supplemented with 100 µg/ml adenine. The following day, cells were diluted 1/10 and grown for 2 h at 25°C. Next, cells were incubated at restrictive temperature (37°C) for 3 h and finally transfer to 384-well CellCarrier plates (PerkinElmer, 6007550) for imaging on an Opera QEHS high-content screening microscope (PerkinElmer). Single planes of 5 different fields were taken per each strain of the library using a 60x water immersion objective and CCD cameras with the proper filter sets to visualise either GFP or RFP. The exposure time for each channel was 1 second. Data analysis to determine the presence of GFP-tagged proteins on Dpb11-labeled anaphase bridges was performed manually. Foci number was also determined by eye.

## ACKNOWLEDGMENTS

We thank other members of the lab for fruitful discussions and help.

This work was supported by the research grants BFU2015-63902-R and BFU2017-83954-R to FM, the Danish Council for Independent Research, the Carlsberg Foundation and the Villum Foundation to ML and OQ, and Canadian Institutes of Health Research grant FDN-159913 to GWB. High-throughput imaging was performed at the Center for Advanced Bioimaging, University of Copenhagen. FM grants were funded by the Spanish Ministry of Economy and Competitiveness (MINECO). Agencia Canaria de Investigación, Innovación y Sociedad de la Información supported CR and SSS through predoctoral fellowships (TESIS20120109 and TESIS2018010034, respectively). All these programs were co-financed with the European Commission’s ERDF structural funds.

## AUTHOR CONTRIBUTION STATEMENT

J.A-P., C.R-P., M.L., G.W.B. and F.M. conceived and designed the experiments. J.A-P., C.R-P., S.S-S., O.Q., S.M-S. and M.Z-D. made the strains and performed the experiments. J.A-P., C.R-P., O.Q., E.M-P. and F.M. prepared the figures and tables. F.M. wrote the manuscript.

## AVAILABILITY OF DATA

The datasets supporting the conclusions of this article are included within the article and its additional files.

