## Supplemental Figures and Tables for "Topoisomerase II deficiency leads to a postreplicative structural shift in all *Saccharomyces cerevisiae* chromosomes"

SUPPLEMENTAL MATERIAL

**SUPPLEMENTAL FIGURES.**

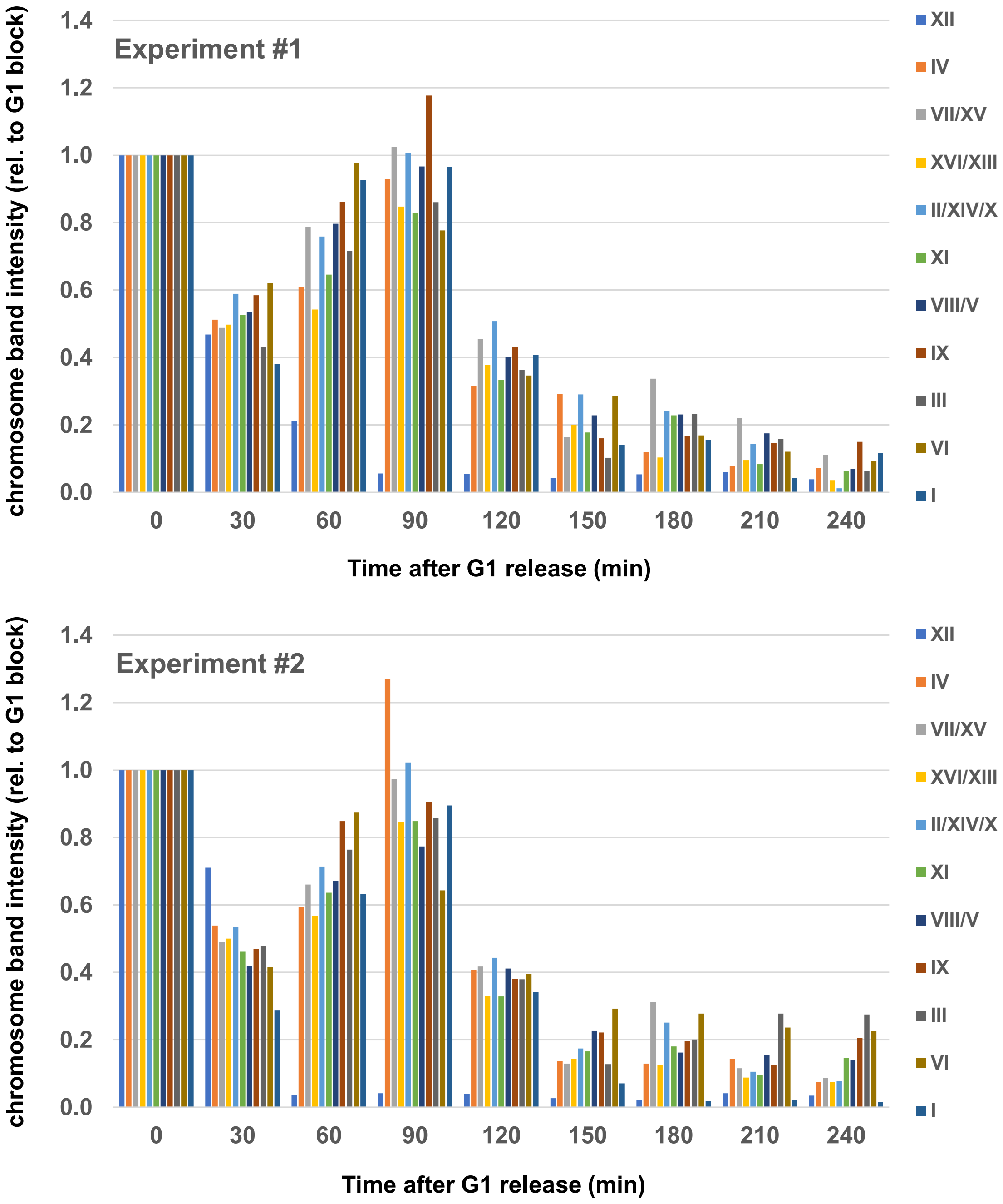


**Figure S1. Quantification of PFG chromosome bands after the *top2-5* release from G1.** After the PFGEs obtained from time course experiments as shown in Figure 1, gels were stained with ethidium bromide, photographed, each band quantified by Fiji/ImageJ (https://imagej.net/Fiji), and represented against the band value at G1. The experiment #1 corresponds to the PFGE shown in Figure 1e.

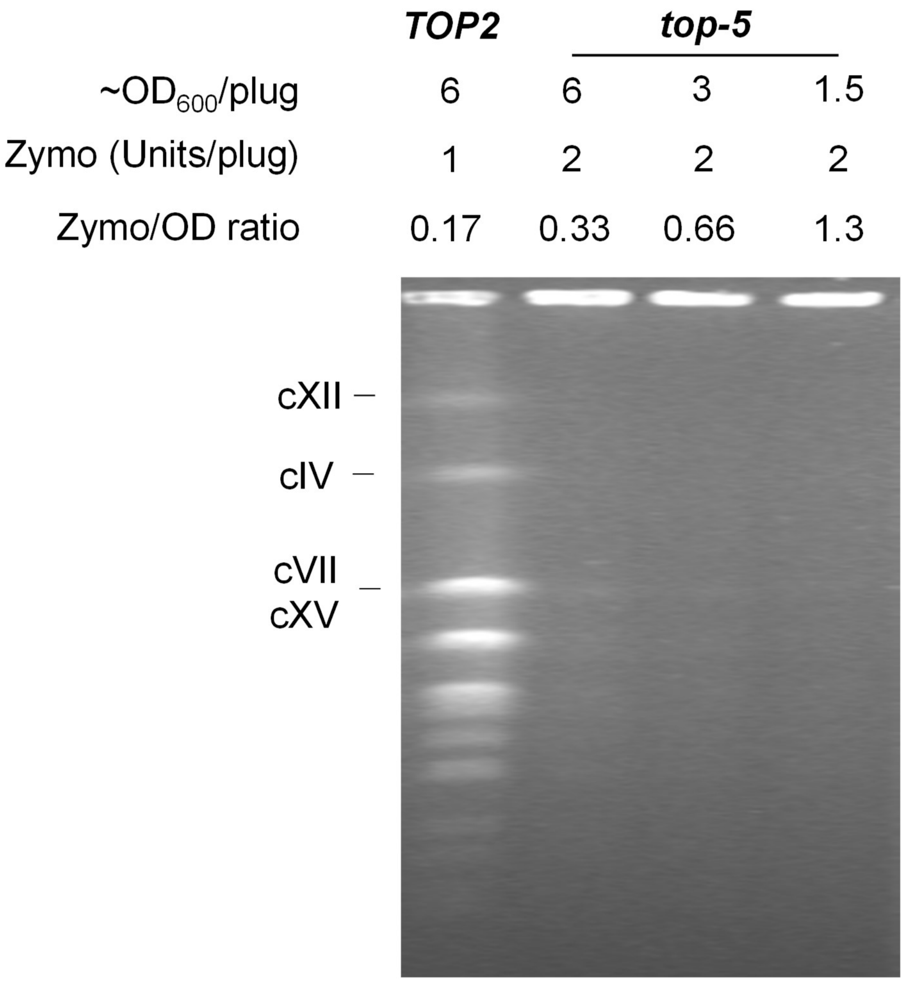


**Figure S2. Well entrapment of *top2-5* is not due to special wall digestion requirements in this mutant.** A time course experiment was carried out as stated in Figure 1. Four hours after the G1 release, samples from the *TOP2* and *top2-5* strains were taken from plug preparation under different cell wall digestion treatments. In general, the number of cells ($OD_{600}$ equivalents) and the amount of zymolyase were changed. The standard values in all other plug preparations of this article were 4 $OD_{600}$ and 2 units zymolyase. Note how increasing ODs and decreasing zymolyase unit in *TOP2* plugs still allowed chromosome migration, whereas decreasing ODs in *top2-5* did not ameliorate chromosome entrapment. The image of the PFG is vertically squeezed to half of its actual length.

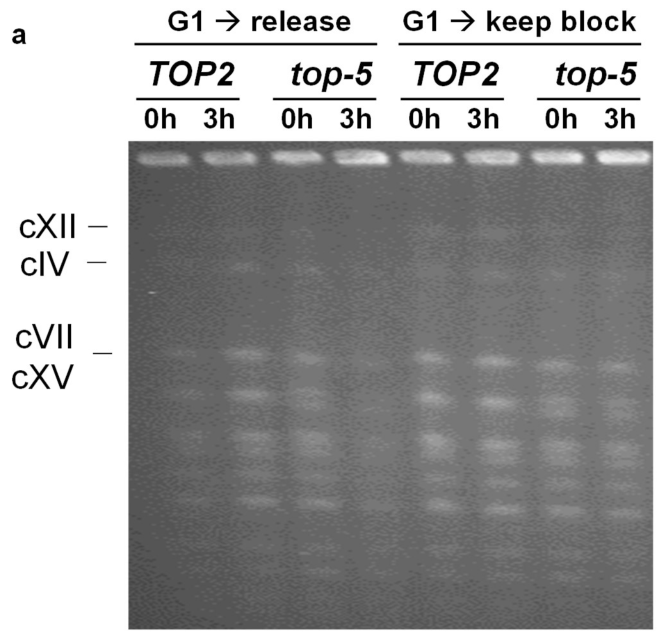


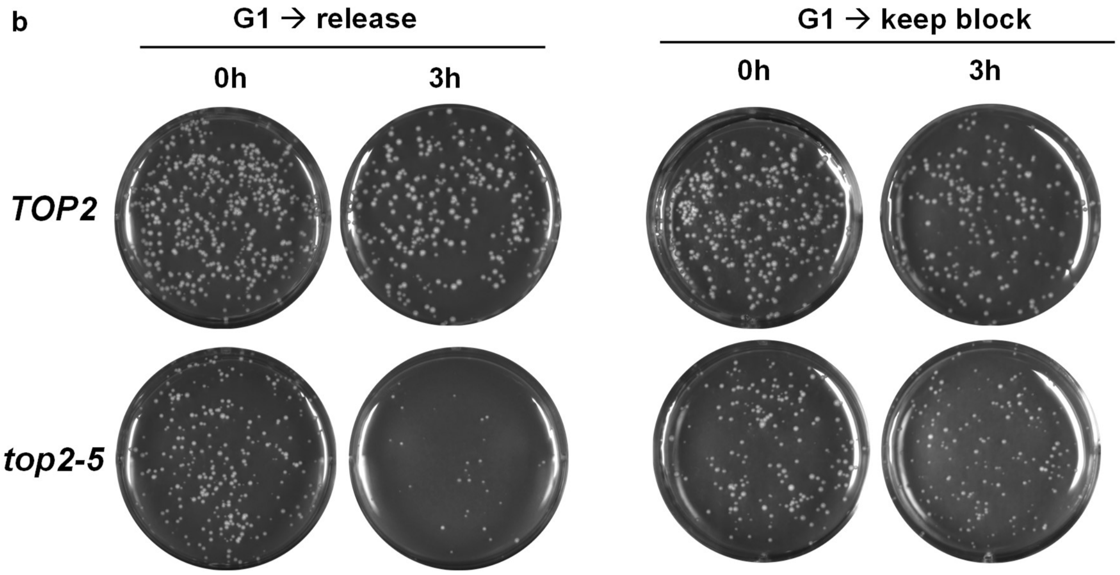


**Figure S3. Maintaining the G1 block prevents chromosomes from well-entrapment in *top2-5*.** The *TOP2* and *top2-5* strains were arrested in G1 for 3h at 25 ºC. Then the cultures were split in two, one of them was released into a synchronous cycle (G1 → release) and the second one was kept in the G1 block. All four cultures were incubated at 37 ºC for 3 more hours before stopping the experiment. At the time of the G1 block at 25 ºC and after 3h at 37 ºC, samples were taken for PFGE and assessment of clonogenic survivability. (**a**) PFGE. The image of the PFG is vertically squeezed to half of its actual length. Note that chromosomes do not get entrapped in *top2-5* cells kept in G1. (**b**) Clonogenic survivability. Note that *top2-5* cells do not loose viability when kept in G1.

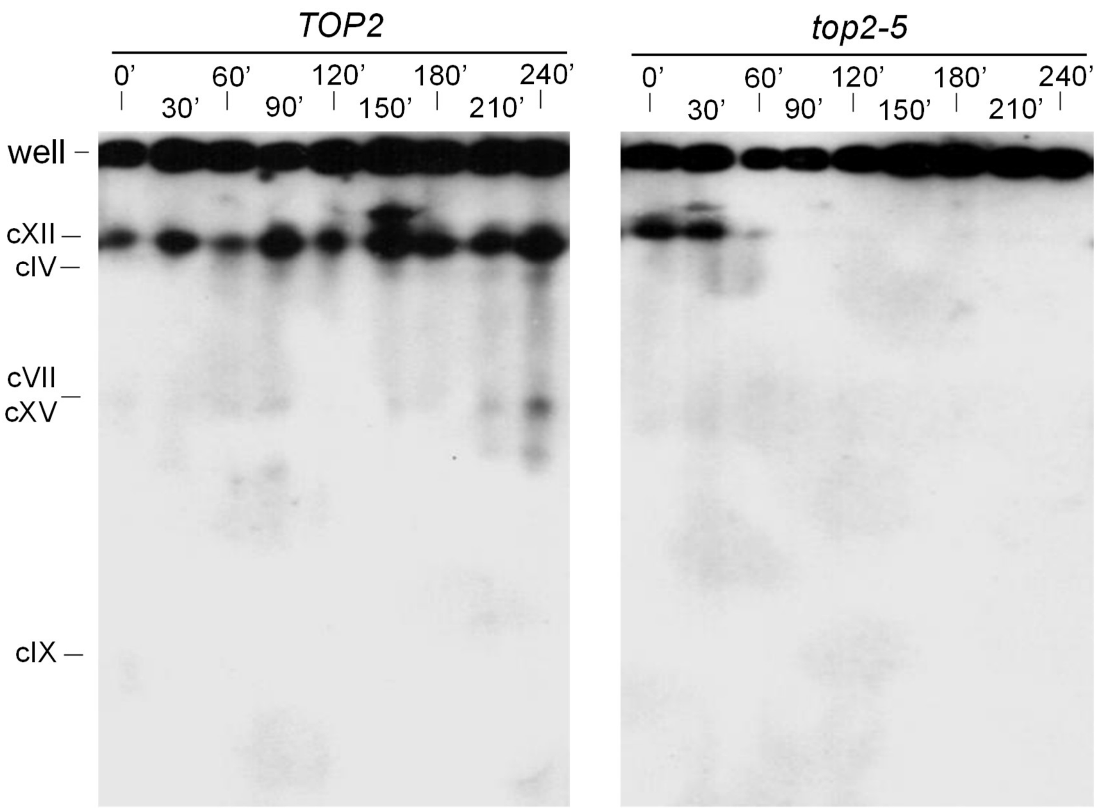


**Figure S4. The mitotic catastrophe upon *top2-5* inactivation entraps chromosome XII in the loading well.** A synchronous G1 release experiment was performed at 37 ºC for isogenic *TOP2* and *top2-5* strains. At the indicated time points, samples were taken for microscopy, FACS and PFGE (not shown). After PFGE, both chromosome bands and well content were transferred to a membrane for a Southern blot analysis with a probe against the NTS1 region of the rDNA (chromosome XII, cXII). Note how no signs of broken cXII (in-gel smear) are observed in *top2-5* despite the observed mitotic catastrophe by FACS (see Figure 1). Lower extra bands were seen for *TOP2* at 210' and 240', but they co-migrated with other chromosome bands. This experiment and the one shown in Figure 1 are independent repetitions.

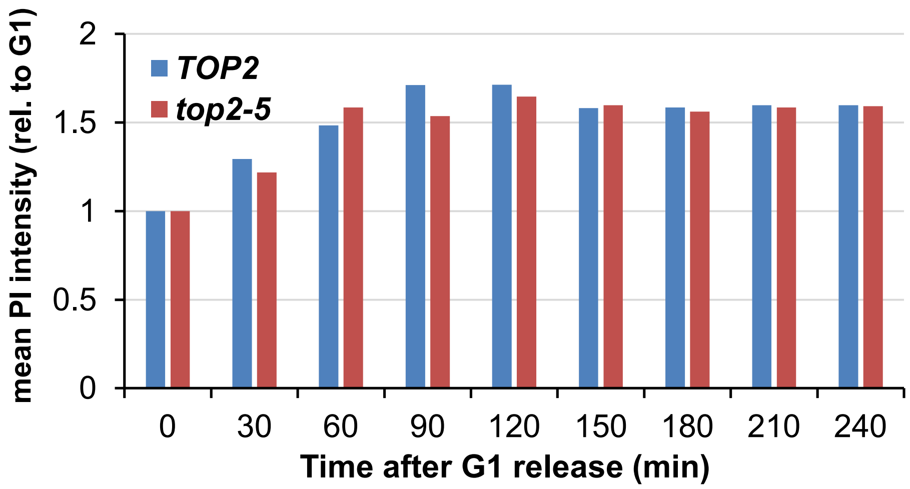


**Figure S5. Quantification of the average DNA content after the *top2-5* G1 release.** Area under the curve was quantified for the FACS profiles shown in Figure 1d. Note that *TOP2* and *top2-5* have an equivalent DNA content during the time course despite the extremely flattened profile of top2-5 at the latest time points (180’-240’). The values never reached the theoretical 2xG1 content, probably because propidium iodine stains budded cells less efficiently.

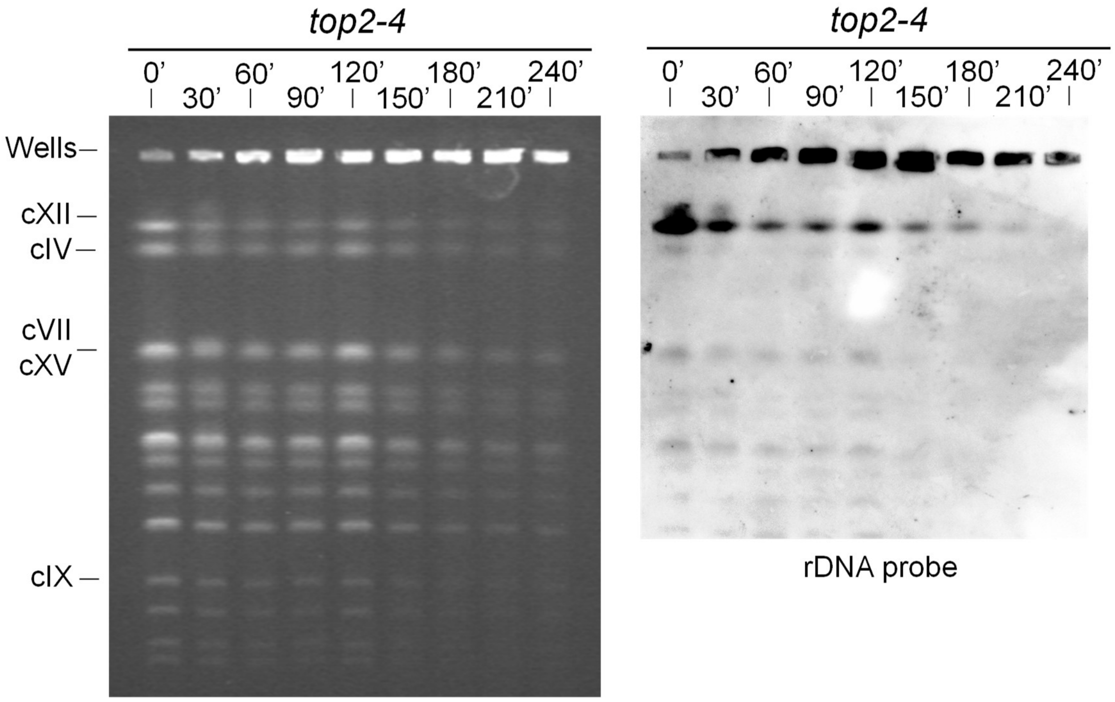


**Figure S6. The mutant *top2-4* also entraps chromosomes after the mitotic catastrophe.** An isogenic strain carrying the *top2-4* ts allele was subjected to the same experiment shown in Figure 1. A PFGE was run and a Southern blot against the rDNA was performed in the same conditions. Note how *top2-4* led to entrapment from 150’ onwards, although the phenotype was slightly milder, including for chromosome XII.

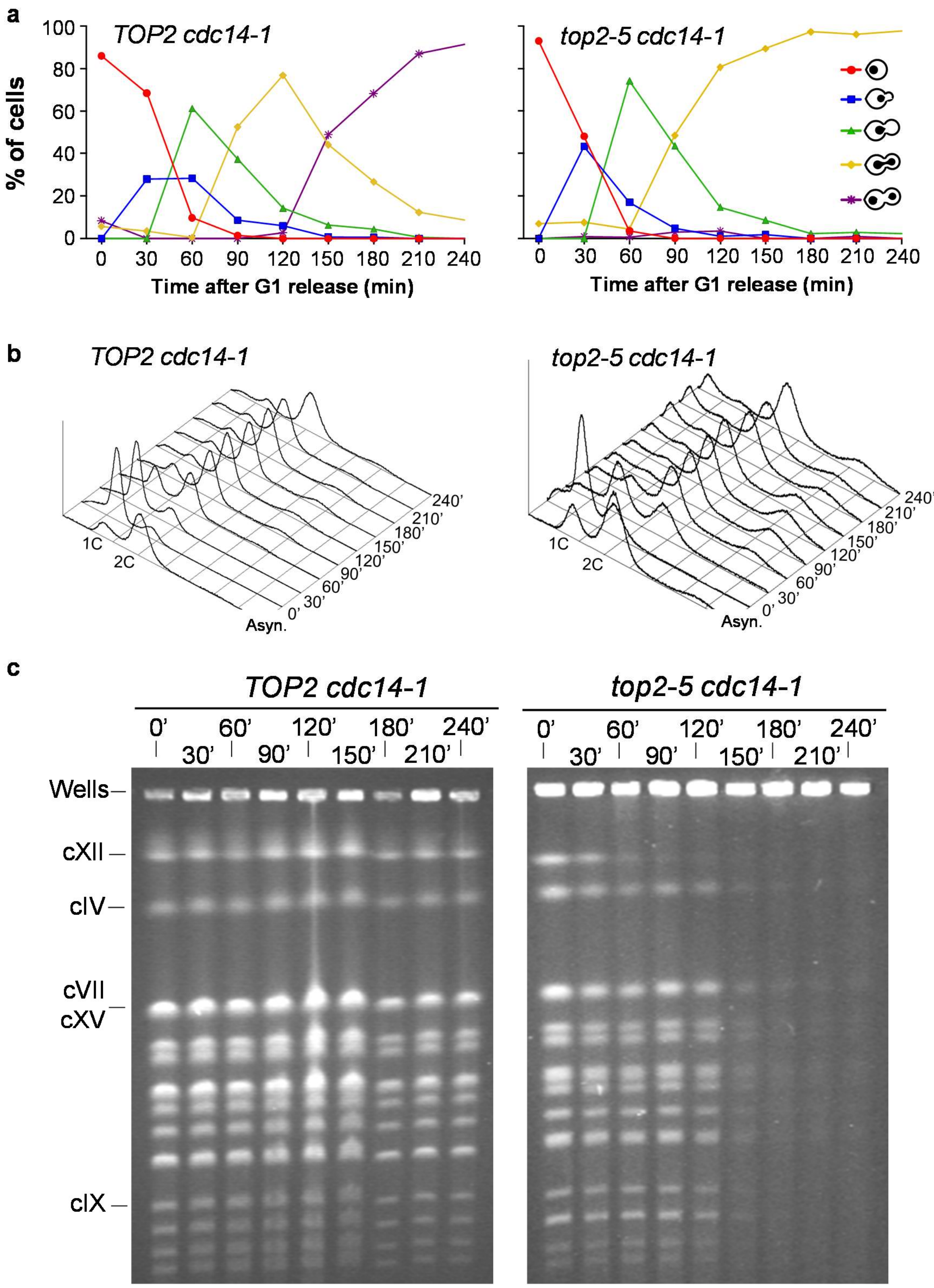


**Figure S7. The loss of chromosome integrity in *top2-5* is not a consequence of Cdc14-driven activities in anaphase.** A synchronous G1 release experiment was performed for isogenic *TOP2 cdc14-1* (left panels) and *top2-5 cdc14-1* (right panels) strains. (**a**) Charts depicting the cell cycle progression under the microscope. (**b**) FACS analysis of the DNA content. (**c**) EtBr staining of whole chromosomes resolved by PFGE. Note how chromosome behaviour of the *top2-5 cdc14-1* strain fully resembled that of *top2-5 cdc15-2*.

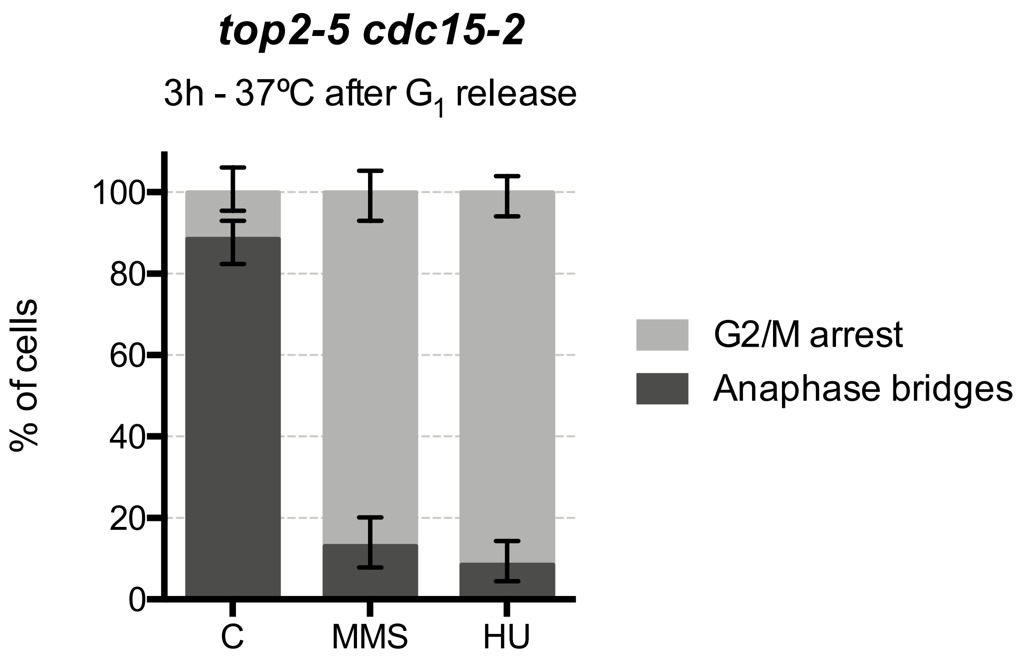


**Figure S8. The replication stress checkpoints are functional in the absence of Top2.** Synchronous G1 release experiments were performed for the *top2-5 cdc15-2* strain. At the time of the G1 release into 37 ºC, the culture was split in three. 2M Hydroxyurea (HU) was added to one subculture; 0.01 % w/v MMS to another one; and the third one was incubated without exogenous replication stress (C). Three hours after the G1 release, samples were observed under the microscope and classify as G2/M (mononucleated dumbbells) or with anaphase bridges (H2A2-GFP stretched across the dumbbell's neck). Error bars depict 95% confidence intervals.

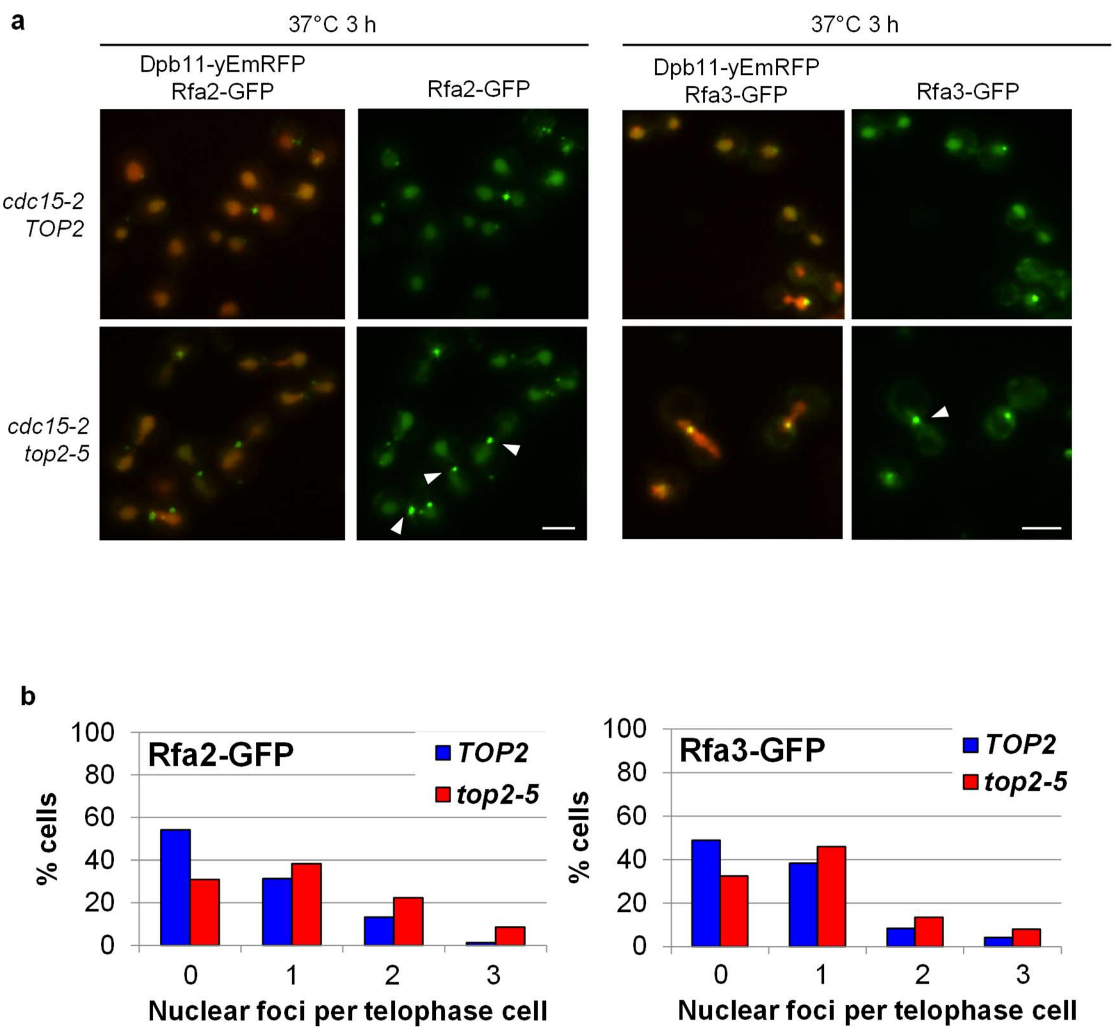


**Figure S9. There is more ssDNA in the *top2-5 cdc15-2* telophase block.** *TOP2* and *top2-5* strains bearing *cdc15-2*, the ultrafine anaphase bridge marker Dpb11-yEmRFP and one RPA component labelled with GFP (either Rfa2 or Rfa3) were blocked in telophase at 37 ºC. (**a**) Representative micrographs. Note how *top2-5* has more foci and they are more intense. Scale bars depict 5 µm. Arrowheads point to foci located at anaphase bridges. (**b**) Quantification of foci number per telophase cell (N>200). See also Table 1.

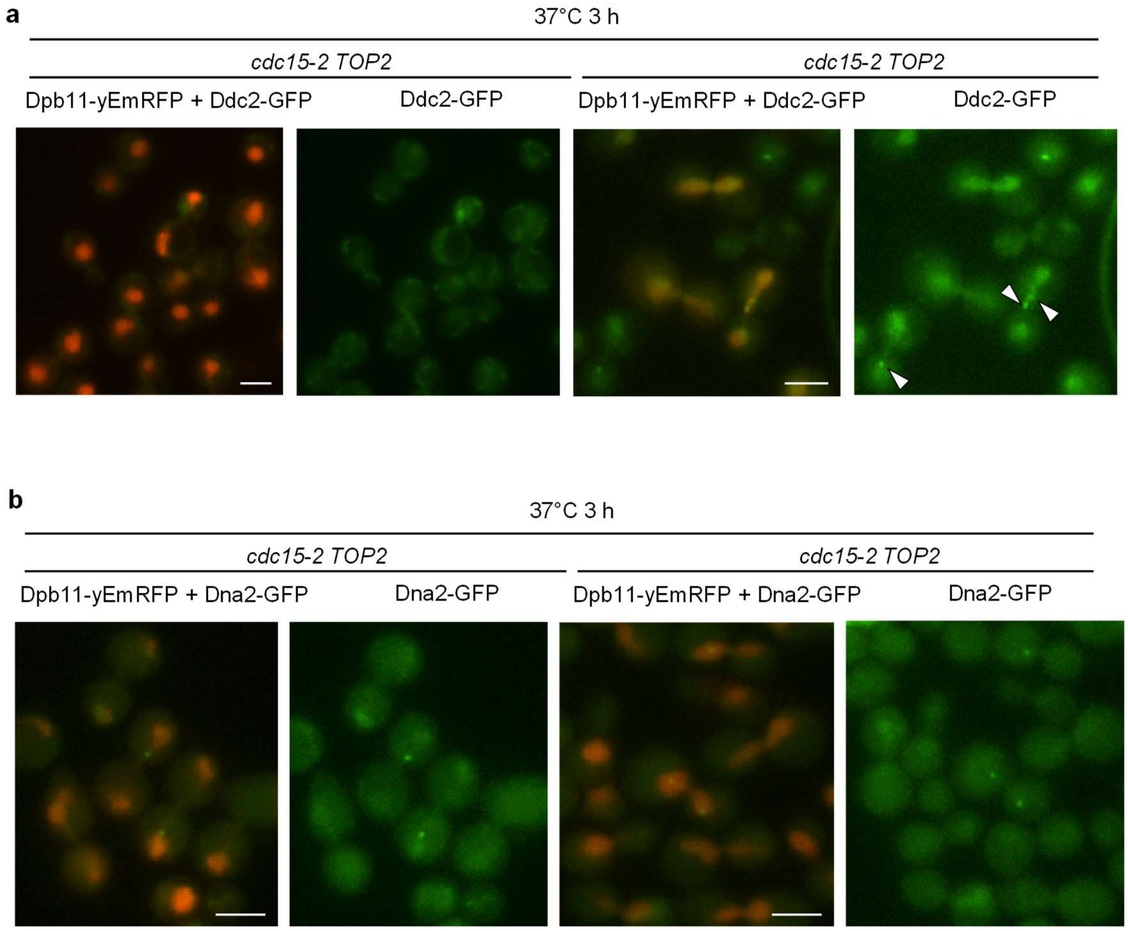


**Figure S10. The replication stress marker Ddc2 gets enriched in *top2-5 cdc15-2* telophase nuclei.** (**a**) Representative micrograph of the *TOP2 cdc15-2 DPB11-yEmRFP DDC2-GFP* and *top2-5 cdc15-2 DPB11-yEmRFP DDC2-GFP* strains blocked in in telophase at 37 ºC. Note that nuclear abundance of Ddc2 increases in *top2-5*, where it also forms foci. (**b**) Representative micrograph of the *TOP2 cdc15-2 DPB11-yEmRFP DNA2-GFP* and *top2-5 cdc15-2 DPB11-yEmRFP DNA2-GFP* strains blocked in in telophase at 37 ºC. Note that Dna2 forms foci in both telophase blocks, regardless the Top2 activity. Scale bars depict 5 µm. Arrowheads point to foci located at anaphase bridges. See also Table 1.

### SUPPLEMENTAL TABLES.

**Table S1. Strains used in this work.**

| Strain name | Relevant genotype[a] | Origin |
|---|---|---|
| CH326 | (S288C) *MAT***a** *ura3-52 his4-539am lys2-801am SUC2+ top2-5* | [1] |
| CH325 | (S288C) *MAT***a** *ura3-52 his4-539am lys2-801am SUC2+ top2-4* | [1] |
| CH335 | (S288C) *MAT***a** *ura3-52 his4-539am lys2-801am SUC2+ TOP2* | [1] |
| FM1386 | CH326; *H2A2(YBL003c):GFP:BleMX*; *Δbar1::URA3* | [2] |
| FM1387 | CH325; *H2A2(YBL003c):GFP:BleMX*; *Δbar1::URA3* | [2] |
| FM1419 | CH335; *H2A2(YBL003c):GFP:BleMX*; *Δbar1::URA3* | [2] |
| FM2021 | FM1386; *cdc15-2:9myc:Hph* | [2] |
| FM2086 | FM1419; *cdc15-2:9myc:Hph* | [2] |
| FM2154 | FM1386; *cdc14-1:9myc:Hph* | [2] |
| FM2152 | FM1419; *cdc14-1:9myc:Hph* | [2] |
| FM2268 | FM2021; *Δrad52::KanMX4* | This work |
| E1670 | (W303) *MAT***a** *ade2-1 trp1-1 can1-100 his3-11,15 leu2- 3,112 RAD5 + GAL psi+ura3::URA3/GPD-*$TK_{7x}$ | [3] |
| FM2128 | E1670; *cdc15-2:9myc:Hph* | This work[b] |
| FM2129 | E1670; *top2-5:NAT; cdc15-2:9myc:Hph* | This work[b] |
| FM2241 | *MATα can1Δ::STE2pr-Sp_his5 lyp1Δ his3Δ1 leu2Δ0 ura3Δ0 met15Δ0 top2-5::natMX* | [2] |
| ML532 | *MATα can1Δ::STE2pr-LEU2 lyp1Δ his3Δ1 leu2Δ0 ura3Δ0 TRP1 LYS2 MET15 KanMX:tTA(tetR-VP16)-tetO2-DPB11* | [4] |
| OQR84 | ML532; *KanMX:tTA(tetR-VP16)-tetO2-DPB11-yEmRFP; top2-5:NAT; cdc15-2:9myc:Hph* | This work[b,c] |
| OQR-TOP2-Xs-GFP (39 strains) | *MAT***a** *can1Δ::STE2pr-LEU2 lyp1Δ his3Δ1 leu2Δ0 ura3Δ0 MET15 KanMX:tTA(tetR-VP16)-tetO2-DPB11- yEmRFP; TOP2; cdc15-2:9myc:Hph; X-GFP:His3MX* | This work[d] |
| OQR-top2-5-Xs-GFP (39 strains) | *MAT***a** *can1Δ::STE2pr-LEU2 lyp1Δ his3Δ1 leu2Δ0 ura3Δ0 MET15 KanMX:tTA(tetR-VP16)-tetO2-DPB11- yEmRFP; top2-5:NAT; cdc15-2:9myc:Hph; X-GFP:His3MX* | This work[d] |

[a] Semicolons separate independent transformation events during strain construction. Intermediate strains are omitted.

[b] The ts alleles were transferred after PCR amplification from strains where they had been tagged downstream with the corresponding markers according to [5].

[c] The yEmRFP was tagged to DPB11 through an adaptamer-mediated PCR fusion methodology described in [6].

[d] These strains were selected after crossing OQR84 with the GFP tagged collection reported in [7], whose basic genotype is *MAT***a** *his3Δ1 leu2Δ0 met15Δ0 ura3Δ0 X-GFP(S65T):His3MX* (*X* being the gene of interest). Individual genotypes are omitted for the sake of space. The haploids were selected on minimum media plates supplemented with canavanine, thialysine, G418 and hygromycin, and in the absence of histidine, leucine and methionine. Then, the presence of either *TOP2* or *top2-5:NAT* was checked by nourseothricin sensitivity or resistance, respectively.
